# A ‘brain-first’ mouse model of progressive alpha-synuclein pathology via intranasal rotenone administration

**DOI:** 10.1101/2025.10.10.677764

**Authors:** Monika Sharma, Nishant Sharma, Jhanvi Soni, Manasi Uttarkar, Jana Ruda-Kucerova, Tiago F Outeiro, Irena Rektorova, Amit Khairnar

## Abstract

The precise aetiology of Parkinson’s disease (PD) is still poorly understood, but it is thought to arise due to an intricate relationship between genes and the environment. Our study takes a unique approach to understanding the effect of environmental factors on the onset and progression of α-synuclein (aSyn) pathology, a key feature of PD, from the olfactory bulb (OB) to other brain regions. In the present study, we evaluated the time-dependent progression of PD-like pathology by administering rotenone intranasally for 5.5 months in C57BL/6 male mice. We performed olfactory and motor tests and examined the aSyn accumulation, glial cell activation and dopaminergic neurodegeneration after 3, 4 and 5.5 months of rotenone exposure by immunoblotting and immunofluorescence techniques.

We observed a time-dependent progression of aSyn accumulation from the OB to other brain regions, including the mid-brain and cortex. Consistently, we observed a time-dependent behavioural impairment, OB atrophy, progression of aSyn pathology, neuroinflammation and neurodegeneration. Our findings also established a link between distinct astrocyte activation and dopaminergic (DAergic) activity. In conclusion, this chronic and progressive mouse model mimics the brain-first type of progression of PD-like pathology in some PD patients, opening the possibility for testing potential disease-modifying interventions.

**Graphical Abstract:** 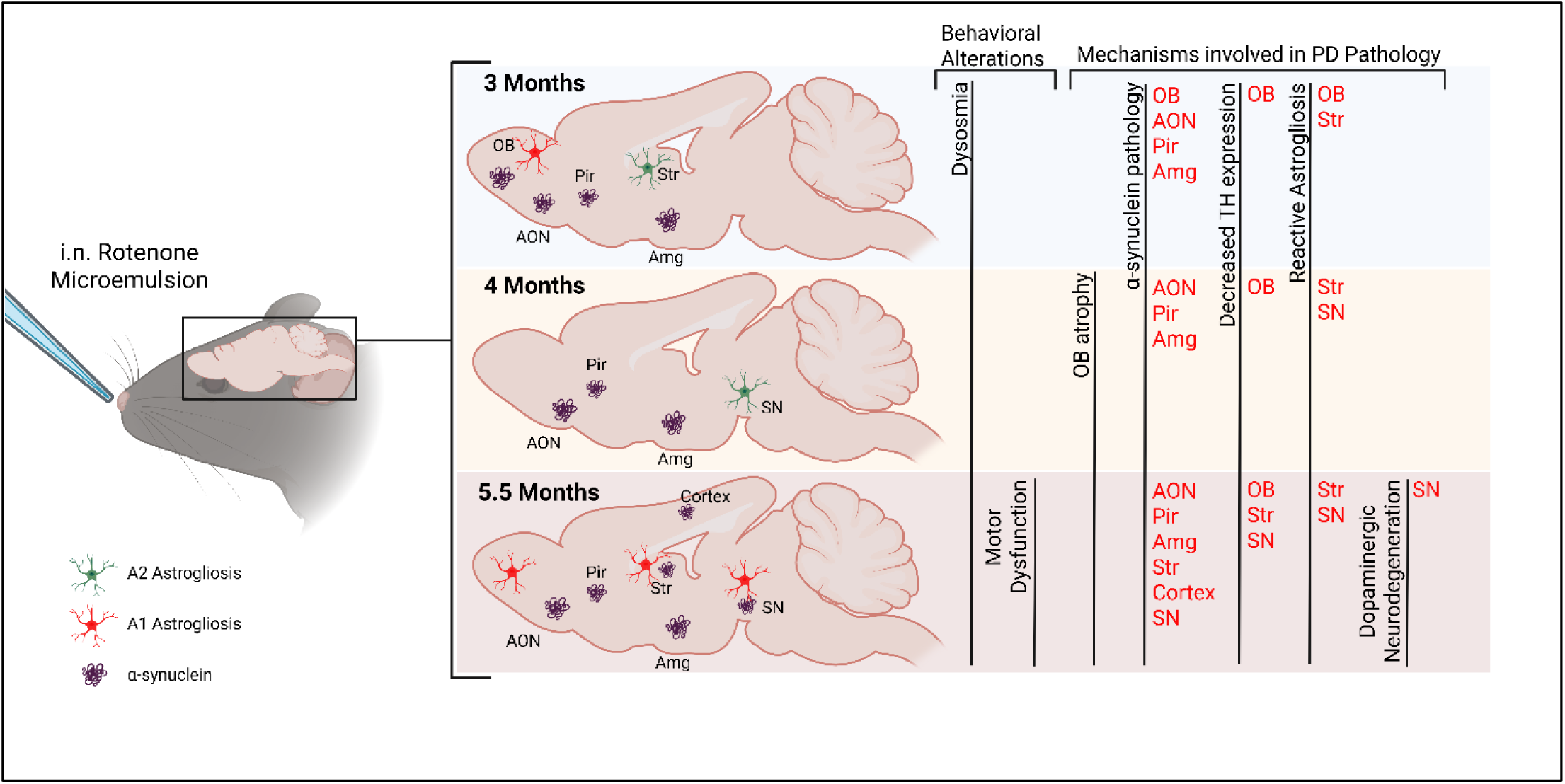

## Introduction

Parkinson’s disease (PD) is a progressive neurodegenerative disorder characterised by substantial loss of dopaminergic neurons in the substantia nigra and depletion of dopamine in the striatum. The accumulation of alpha-synuclein (aSyn) containing inclusions known as Lewy bodies and Lewy neurites is a pathological hallmark of PD and other Lewy body diseases. It is characterised by classic motor symptoms such as bradykinesia (slowness of movement), rigidity, and resting tremor, which usually appear after a long preclinical phase of accumulating pathology and often mark the point at which clinical diagnosis is made. Non-motor symptoms are prevalent throughout the stages of PD and significantly impact patients’ quality of life. These symptoms encompass a broad range of clinical features, leading to considerable morbidity. In humans, aSyn pathology manifests several years before the onset of the classical motor symptoms of PD, and it is frequently preceded by non-motor deficits such as olfactory dysfunction, constipation, and sleep disturbances. (Berg et al. 2015, Attems et al. 2014). Neuronal loss and synucleinopathy have been identified not just in the substantia nigra (SN) but also in other parts of the brain, including but not limited to the olfactory bulb (OB), anterior olfactory nucleus (AON), amygdala, and piriform cortex (Torres-Pasillas et al. 2023). Olfactory dysfunction is an early and often overlooked aspect of PD, characterised by the accumulation of aSyn in the olfactory bulb, a region crucial for the sense of smell, leading to hyposmia, which is prevalent in the majority of patients in earlier stages.

Through the examination of postmortem human brain tissue of individuals diagnosed with PD, Braak hypothesised the spreading of aSyn pathology from the OB and AON to other brain regions, via axonal projections (Braak *et al*. 2003). Studies using a variety of in vitro and in vivo models supported this hypothesis, suggesting aSyn pathology may spread in a prion-like manner. (Li et al. 2008; Kordower et al. 2008; Luk *et al*. 2009; Volpicelli-Daley *et al*. 2011; Hansen *et al*. 2011;Mougenot *et al*. 2012; Bernis *et al*. 2015). Importantly, there are two possible origins of aSyn pathology: the brain-first or the body-first hypothesis. In the body’s first hypothesis, the prodromal phase is more extended, and the patient has positive autonomic dysfunction, such as RBD and constipation.(Horsager & Borghammer, 2024)(Borghammer et al., 2021) Rotenone, a natural pesticide, has been used by our group to develop the body first hypothesis in mice, where the aSyn pathology develops in the gut and then it progresses to the midbrain through the dorsal motor nucleus of vagus (Francisco PM et al., 2010, Khairnar et al., 2021, Sharma N et al., 2023). While the brain in first PD patients bypasses the gut and autonomic dysfunction, showing origin of aSyn pathology either in the amygdala or the olfactory bulb, with the olfactory dysfunction as a primary non-motor symptom, with a lower prodromal phase (Attems *et al*. 2014, Fullard ME et al., 2017, Horsager J et al., 2020). Hence, the current study focused on developing a brain-first hypothesis mouse model using intranasal rotenone that would provide a valuable tool for testing innovative disease-modifying treatments, which may halt the pathological process.

In several studies, authors have targeted OB in PD and have tried to expose the nasal mucosa to neurotoxicants, which might be responsible for inducing PD pathology in the brain via the olfactory mucosa. These studies include intranasal administration of dopaminergic toxins or pesticides (MPTP, 6-OHDA, rotenone and paraquat) in rodents, leading to PD-like pathology. (Prediger *et al*. 2012; Rojo *et al*. 2007; Sasajima *et al*. 2015; Sasajima *et al*. 2017; Kawano & Margolis 1982). Targeting the OB as the initiation site of aSyn pathology was also reported, revealing the rapid uptake of human recombinant aSyn by the OB and its axonal transit to many interconnected brain areas after retrobulbar injection in mice (Rey *et al*. 2013; Rey *et al*. 2016; Rey *et al*. 2018a). Although informative as experimental tools, these models are very acute, as this situation never happens in humans. Therefore, it is important to continue developing models that may more faithfully recapitulate sporadic forms of PD, or forms that depend on environmental factors, such as exposure to pesticides.

Rotenone is commonly used to induce PD-like phenotypes in animal models (Sharma M *et al*. 2023, Bandookwala M *et al*. 2019), and it triggers the buildup of aSyn pathology in the intestine (Drolet *et al*. 2009; Ishola *et al*. 2023) or nasal cavity (Voronkov *et al*. 2017; Prediger *et al*. 2012). Therefore, following the Braak hypothesis, interneurons extending from the olfactory bulb to the nasal mucosa exposed to rotenone should lead to aSyn accumulation in the OB, with subsequent spreading to other parts of the brain.

Here, we intranasally delivered a locally-acting, low-dose rotenone microemulsion (ME) in adult mice that does not exert measurable concentrations in the blood, OB, or other brain regions (Sharma et al. 2022). We administered rotenone ME chronically and evaluated behavioural parameters and the progression of aSyn pathology in numerous brain regions (OB, AON, piriform cortex, amygdala, striatum, SN, and cortex) at several time points. Moreover, we investigated additional pathological markers, including neuroinflammation, DAergic activity, and neurodegeneration in the OB, striatum, SN, and cortex. Lastly, we examined a probable mechanism underlying the distinct time-dependent astroglial pathology in different brain regions and its relation to DAergic activity.

## Materials and methods

### Animals

Three-month-old C57BL/6 mice (a total of 60 mice) were procured from Zydus Research Centre, Ahmedabad, after being approved by the Institutional Animal Ethics Committee (Approval number: IAEC/2021/010) of NIPER-Ahmedabad. Animals were kept at proper temperature (18-23°C) and moisture conditions (40-60% humidity) under a 12 h light/dark cycle (with lights switched off during the dark cycle) and provided with food and water *ad libitum*. All the experiments were conducted per the guidelines of the Committee for the Purpose of Control and Supervision of Experiments on Animals (CPCSEA), India.

### Chemicals

Rotenone, acrylamide, sodium dodecyl sulfate (SDS), tween-20, formaldehyde, 3,3′-diaminobenzidine (DAB), glucose oxidase, and D-glucose were obtained from Sigma Aldrich. Ammonium persulfate (APS), tetramethyl-ethylenediamine (TEMED), sodium chloride, sodium lauryl sulfate (SLS), bis-acrylamide, sodium deoxycholate, Triton-X 100, phenylmethyl sulfonyl fluoride (PMSF), glycerol, and carboxymethylcellulose (CMC) were purchased from Hi-Media Laboratories Pvt. Ltd. β-mercaptoethanol was purchased from Alfa Aesar. Potassium chloride, hydrochloric acid, sodium hydroxide, and chloroform were procured from Fischer Scientific. Polyethene glycol-400 was purchased from Merck. Tris-HCl was purchased from Thermo Fischer. The protein ladder and PVDF membrane were procured from Bio-Rad. HRP conjugated Enhanced Chemiluminescent Substrate Reagent kit was obtained from Invitrogen. The BCA (bicinchoninic acid) reagent kit was bought from Thermo Scientific. Primary antibodies (GAPDH, aSyn, phosphorylated aSyn (psyn), S100A10, GDNF) and secondary HRP-conjugated and secondary fluorescent antibodies were purchased from Abcam. Primary antibodies: glial fibrillary acidic protein (GFAP, AB5804), tyrosine hydroxylase (TH, AB152) and phosphorylated TH (pTH, AB5935) were procured from Sigma-Aldrich. Complement C3 was obtained from Thermo Scientific. For immunohistochemical staining, the Vectastain ABC kit was procured from Vector Laboratories.

### Study plan

Rotenone mucoadhesive ME (0.1 mg/kg) was prepared as described in our previous study (Sharma *et al*. 2022). The same microemulsion without rotenone was used as the vehicle. The animals were divided into a vehicle-treated control group and a rotenone ME-treated group, n=10 for each of the three time points: 3, 4 and 5.5 months. Each mouse was placed on its back for the ME administration and given a 15 µL ME intranasally via micropipette for 5 days/week. After completion of 3, 4 or 5.5 months, animals were sacrificed by cervical dislocation and their brains were quickly removed on ice and OB, striatum and cortex were extracted from one hemisphere and preserved in liquid nitrogen and then stored at -80°C until western blotting. While the other hemisphere was collected in a 25 ml Falcon tube and kept at 4°C in a 4% PFA solution (phosphate buffer saline, PBS, pH 7.4) overnight and later it was transferred to 15% sucrose in PBS for further immunohistochemistry studies (Fig. 1A).

**Figure 1.**
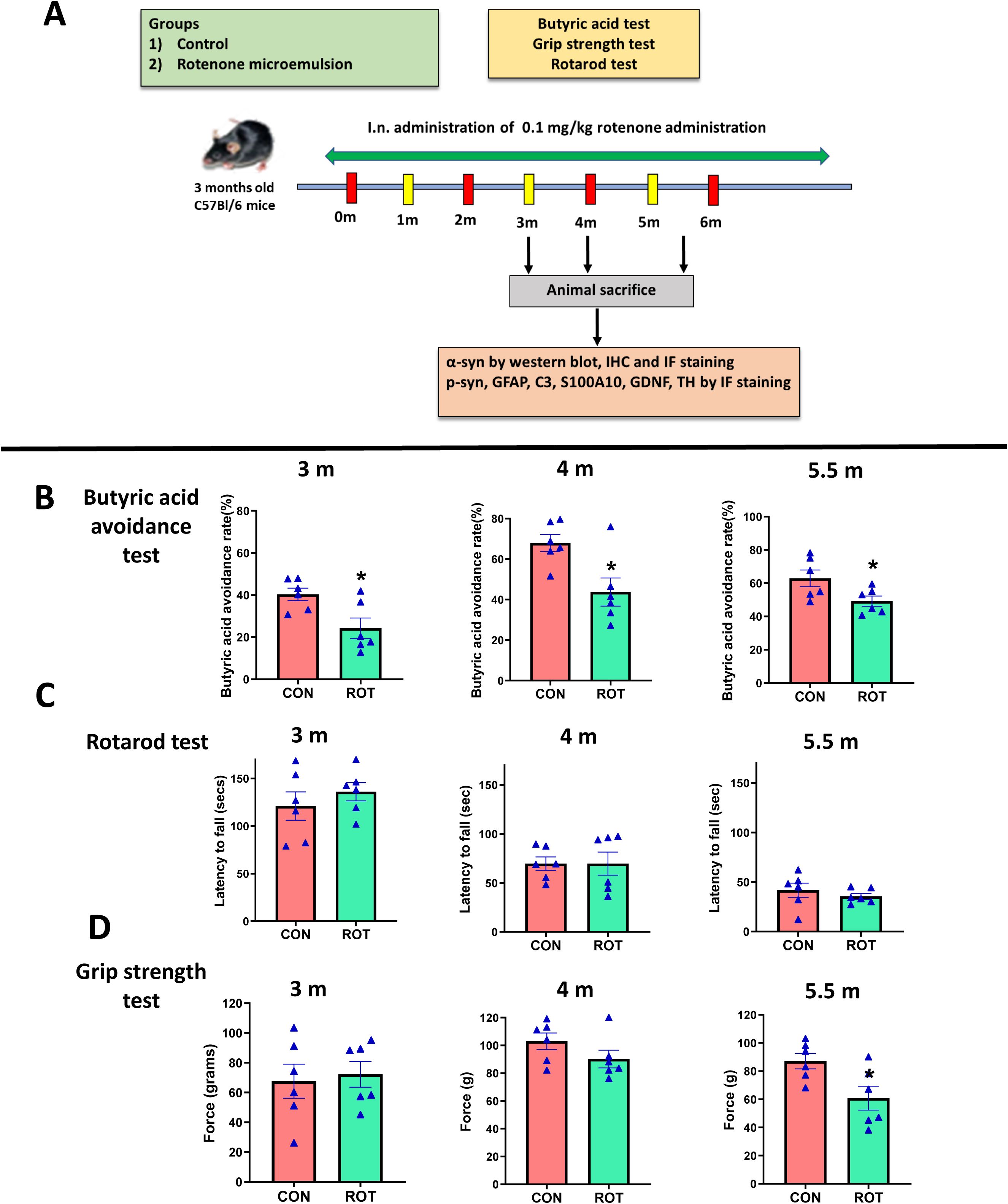
Experimental design used in the study. A) Diagrammatic illustration of the experimental procedure. (B) Rotenone induced olfactory dysfunction after 3, 4 as well as 5.5 months of intranasal administration: Graphs represent the avoidance ratio of butyric acid measured by Y-maze apparatus in control (CON) and rotenone (ROT) animals after 3 months, 4 months and 5.5 months (m) of rotenone administration. Data was analyzed by two-way ANOVA followed by post-hoc ” ‘Tukey’s test and expressed as mean ± SEM (n=6 in both control and rotenone group), *p<0.05. (C) Rotenone induced motor dysfunction after 5.5 months of intranasal administration: Graphs represent the force applied by mouse paws in grip strength test in control and rotenone treated animals after 3 months, 4 months and 5.5 months of i.n. rotenone ME administration respectively. Data was analyzed by unpaired t-test and expressed as mean ± SEM (n=6 in both control and rotenone group), *p<0.05. (D) Rotenone didn’t induce any change in latency to fall after 3, 4 and 5.5 months of intranasal administration: Graphs represent the latency to fall from the rod in rotarod test in control and rotenone animals after 3 months, 4 months and 5.5 months of i.n. rotenone ME administration. Data was analyzed by unpaired t-test and expressed as mean ± SEM (n=6 in both control and rotenone group).

### Behavioral tests

The following behavioural tests were conducted to check the olfactory deficit and motor and memory impairment at all time points before sacrificing the animals.

#### Butyric acid avoidance test (BAT)

This test was conducted using a Y-maze as described by Sasajima et al., with minor modifications (Sasajima et al. 2017). Two filter papers saturated with 20µL of water in petri dish were kept in two arms of the Y-maze, and the animal was placed at the end of the third arm and permitted to explore the apparatus for 4 minutes. On the following day of the experiment, water was replaced with the same volume of butyric acid in one arm, and the animal’s time spent with the petri plates in both arms was recorded for 4 minutes. Then, the butyric acid avoidance ratio was calculated as (time spent in arm with water)/(time spent in arm with butyric acid + time spent in arm with water).

#### Grip strength test

The muscle strength of mice was tested using a grip strength meter. The mouse was placed on the grid and moved on it with all four paws while holding onto its tail. A grip strength meter was used to measure the maximal force (g) applied by mouse paws to hold the grid. This was done in triplicate for each mouse, with a 30-minute interval between trials (Sharma *et al*. 2022). The average of three trials was used for statistical analysis.

#### Rotarod test

Motor coordination of mice was assessed using the rotarod apparatus (Harvard apparatus) (Shiotsuki *et al*. 2010). All animals were trained to walk on the rotating rod at a constant speed of 4 rpm for three days. After training, the final test was carried out in an acceleration mode (2 to 20 rpm for 300 seconds). Each animal was tested three times with 30-minute intervals, and the time the animal took to fall from the rod was recorded. The average of three trials was used for statistical analysis.

### Reverse transcriptase polymerase chain reaction (qrt-PCR)

Total RNA was isolated from the cells using the TRIZOL method per the manufacturer’s instructions, and purity was quantified using a Nanodrop 2000c spectrophotometer (Thermo Scientific, Wilmington, DE). Extracted RNA was reverse transcribed using cDNA synthesis kit (Bio-Rad Laboratories, USA). The prepared cDNA was diluted using SYBR® Green Supermix (Bio-Rad, USA) and primers with Bio-Rad CFX 96™ Real-Time system (Bio-Rad, USA). The quantification of interleukin-1 beta (Il-1β), interleukin-6 (IL-6) and interleukin-10 (IL-10) genes was carried out by the ΔΔCT method with 18s as endogenous control. The primer sequence for evaluated markers is mentioned in Table 1.

**Table 1.**
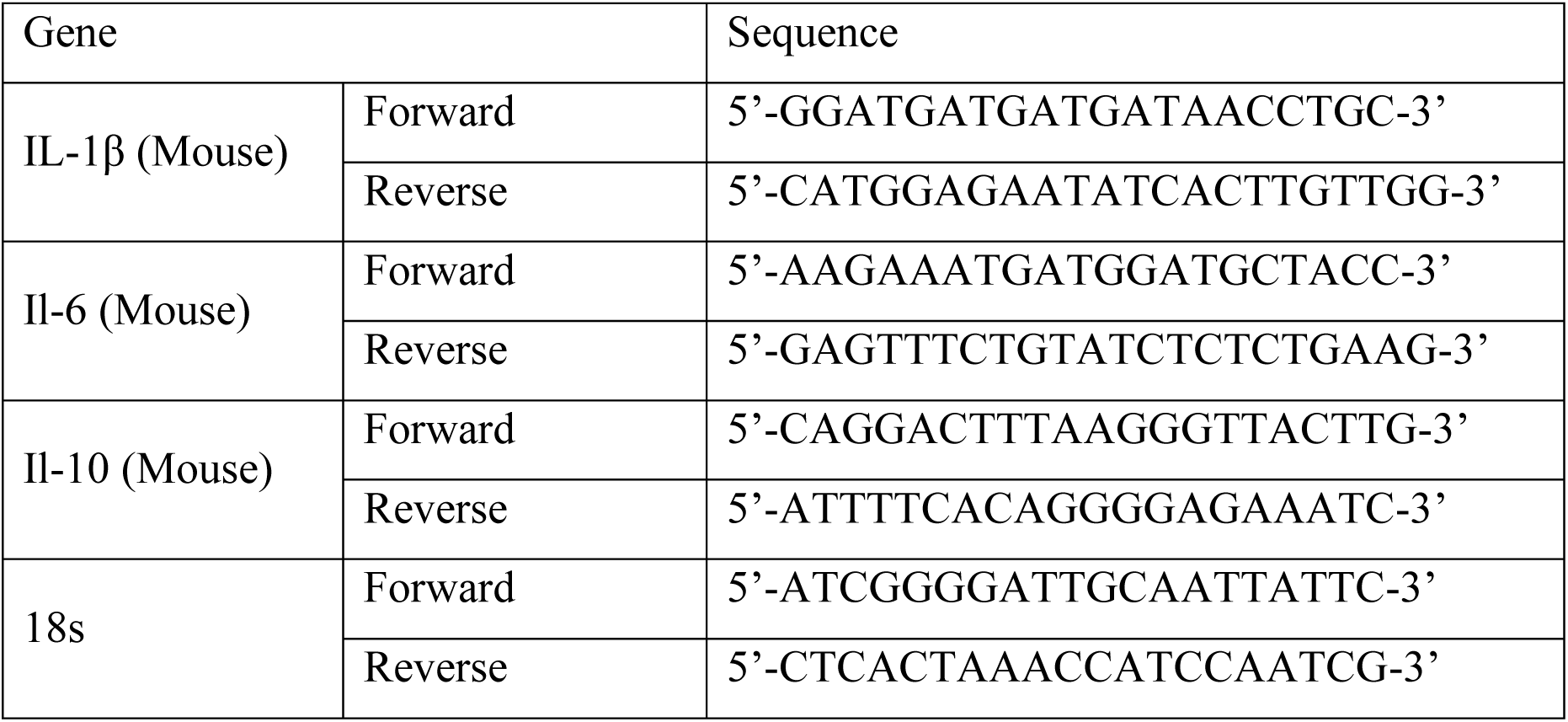
Sequences of primers used in this study.

### Western blotting (WB)

Frozen OB, striatum and cortex samples were thawed, and tissue lysates were prepared using RIPA buffer, which consisted of Tris-HCl, NaCl, SDS, Sodium deoxycholate and 1% Triton X-100 mixed with PMSF (protease inhibitor). Lysate samples were centrifuged at 12,000 rpm for 7 minutes, and supernatants were collected and used for analysis. The Bicinchoninic acid (BCA) technique was used to determine protein concentration. Thirty µg of protein was separated on 15% and 12% SDS-PAGE gels and transferred to the PDVF membrane using the Bio-Rad Trans-Blot assembly. Membranes were then blocked using 3% BSA and incubated with primary antibodies of GAPDH (ab8245, 1:10000), aSyn (rabbit pAb, ab212184, 1:1000), phosphorylated aSyn (rabbit mAb, ab51253, 1:1000), tyrosine hydroxylase (rabbit pAb, AB152, 1:10,000), phosphorylated tyrosine hydroxylase ((pTH) (rabbit pAb, AB5935, 1:1000)) GFAP (rabbit pAb, AB5804, 1:1000), S100A10 (rabbit mAb, ab76472, 1:1000),

Complement C3 (rabbit pAb, PA5-21349, 1:500) and GDNF (rabbit pAb, ab18956, 1:500), overnight at 4°C with gentle shaking. The next day, antigen-antibody complexed membranes were washed with TBST and further incubated with respective HRP-conjugated secondary antibodies (goat anti-rabbit pAb, ab6721, 1:10000; goat anti-mouse pAb, ab6789, 1:10000; rabbit anti-goat pAb, ab6741, 1:10000), followed by chemiluminescent detection. GAPDH was used as an internal control to normalise the protein levels. The expression level was quantified by densitometric analysis with the help of Image J software (NIH, USA) (Parkhe *et al*. 2020).

### Immunohistochemistry staining (IHC)

Brain hemispheres of one side were stored in 4% PFA overnight and then transferred to 15% sucrose (sucrose solution in 0.1 M phosphate buffer pH 7.4) at 4 °C for 24 hours before being submerged in 30% sucrose solution. After that, these hemispheres were processed for cryostat sectioning. Cryostat (Thermo Fisher Scientific) was used to collect 40 µm free-floating coronal sections of SN of control and rotenone mice at six different levels in 24-well culture plates and processed for IHC-DAB staining for TH. According to our previous study, Immunohistochemistry was performed using the Vectastain ABC kit, with minimal modifications (Khairnar *et al*. 2010). For 10 minutes, sections were incubated in a 1% hydrogen peroxide solution in 0.1 % Triton-X-100 and then blocked with 5% normal goat serum in 0.1% Triton-X-100 in PBS. After that, sections were incubated with anti-TH (rabbit pAb, AB152, 1:1000) antibody overnight at 4°C. The next day, PBS washing was done, followed by secondary biotinylated antibody (for 1^1/2^ h) and avidin-biotin-peroxidase (Diluted ABC solution, Vector Laboratories) for one hour in the dark at room temperature with intermediate three PBS washings of 10 minutes each. The peroxidase reaction was developed with DAB substrate in the presence of glucose and ammonium chloride dissolved in 0.1 M phosphate buffer for 5 minutes, followed by glucose oxidase incubation for 8-15 minutes. Then, sections were taken on slides, dried and dehydrated in ethanol gradient solutions. DPX was used for mounting, and then images were taken using a Leica DMi1 inverted microscope. The same processing was done with sucrose-processed sections of OB, AON, piriform cortex, amygdala and SN after cryosectioning at 40 µm thickness. The primary antibody was aSyn (rabbit pAb, ab212184, 1:1000).

#### Quantitative analysis of DAB staining

A. Dopaminergic neuronal count in SNc: We used a confocal microscope to perform stereological counting. Images were acquired using a confocal microscope (Leica TCS SP8 Microsystem). As defined by Ip et al. (Ip *et al*. 2017). Six randomly selected sections, separated by 240 µm (1/6 series) along the whole anterior-posterior extent of SNc, were subjected to counting. TH-immunoreactive DAergic neuronal perikarya were recognised by their rounded or ovoid shape. To perform TH stereological counting, we employed the following parameters: counting frame size (50 μm × 50 μm), and sampling grid size (130 µm × 130 µm). The following formula was used to compute the estimated number of TH^+^ positive cells per animal (N).

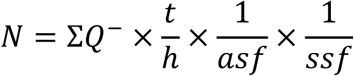

Where ƩQ is the sum of all TH^+^ neurons counted in all optical dissectors of a single brain section; h is the height of the optical dissector, t is the mean tissue thickness of the section; asf is the area sampling fraction defined as the proportion of the area (A) of the optical dissector frame size within the square size of grid (A x,y step); ssf is the section sampling fraction defined as the proportion of sections of the whole serially cut brain. The final N values were only included if their coefficient of error (CE) was less than 10. The researchers did the analysis blinded to the experimental groups.

#### B. aSyn intensity in OB, AON, piriform cortex, amygdala

Image analysis was done using the ImageJ software (National Institutes of Health (NIH), Bethesda, MD, USA) as described in the previous literature (Jewett *et al*. 2017; Wang *et al*. 2018; Xavier *et al*. 2005). The extent of immunostaining of aSyn in OB, AON, piriform cortex, and amygdala was presented as a percentage of optical density in control mice. Image J was first calibrated using the Rodbard function within the software to normalise the grey-scale range (0–255) into OD values. Each image was transformed into an 8-bit (grey-scale) image. The OD values were then normalised by eliminating the OD values of the background and normalising them to control values.

### Immunofluorescence staining and analysis

Free-floating coronal sections (40µm) of SN at three different levels (-3.52mm, -3.16mm, - 2.80 mm) of control and rotenone mice brains were processed for immunofluorescence staining. The sections were rinsed three times with 0.1 M PB before being blocked for 20 minutes with protein block (ab64226). Primary antibodies were diluted in Antibody diluent (ab64211) and incubated with the sections overnight at 4°C. The sections were then rinsed three times in PB containing 0.025% triton-x before being incubated for one hour at room temperature with secondary antibodies as per our previous studies (Sharma M *et al*. 2022, Khairnar *et al*. 2016). Primary antibodies used are anti-TH (rabbit pAb, AB152, 1:500), anti aSyn (mouse mAb, AB1903, 1:500), anti-GFAP (rabbit pAb, AB7260, 1:500) and, S100A10 (rabbit mAb, ab76472, 1:100), Complement C3 (rabbit pAb, PA5-21349, 1:200) and GDNF (rabbit pAb, ab18956, 1:50). Secondary antibodies used are goat pAb secondary to mouse, Alexa-Fluor 488 (ab150113) (1:1000), and goat pAb secondary to rabbit, Alexa-Fluor 647 (ab150079) (1:1000) and donkey pAb secondary to goat, Alexa-Fluor 555 (ab150134).

The nuclei were stained with DAPI after another rinse with PB mixed with Triton X (Sigma-Aldrich, USA). Images were acquired using a confocal scanning laser microscope (Leica TCS SP8 Microsystem). Using ImageJ software (version 1.42, NIH, USA), mean fluorescence from aSyn, GFAP, C3, S100A10, and GDNF was evaluated for all three levels in SNc (Farrand *et al*. 2020). For each region, background measurements were obtained and removed from the mean fluorescence levels, with the corrected values shown as a percentage of the control.

### Nissl staining and image analysis

To check the atrophy in the OB, we stained 40 µm coronal sections of the OB with Nissl stain. Free-floating sections were taken on slides and air-dried. Then, sections were rehydrated by dipping them into 90%, 80%, and 70% ethanol, then rinsing the slide in water. Subsequently, sections were incubated with 0.1% cresyl violet (warm it at 60^ο^C for 10-15 min) for 15 min. After this, sections were dehydrated using alcohol gradients, followed by xylene treatment. Then, the sections were mounted using DPX, and images were captured using a Leica DMi1 inverted microscope (Tepper *et al*. 2021). The area of different layers of OB was analysed using ImageJ software. First, the image was opened in the software, accompanied by setting the scale in µm and then the area of interest was drawn using the freehand selection tool, followed by measuring and analysing.

### Statistical analysis

GraphPad Prism, version 5.01, GraphPad Inc. software was used for statistical analysis. All the data were expressed as mean ± SEM. All data passed the Shapiro-Wilk test of normality. A two-sided Student’s t-test was used to compare control and rotenone groups, except for the analysis of BAT, where a two-way ANOVA test was used, followed by post-hoc Tukey’s test. The boundary for statistical significance was set to p<0.05.

## Results

### Rotenone induces time-dependent behavioural impairment

To develop a model of the brain-first subtype of PD, we administered rotenone intranasally and assessed the animals at different time points. We found that there was a significant (p<0.05) increase in butyric acid avoidance ratio in control mice. In contrast, rotenone-administered mice did not show any considerable avoidance at any time point (Fig. 1B). This showed that rotenone administration led to olfactory dysfunction in mice after 3 months of administration. There was no significant difference in force applied by mice paws in grip strength test after 3 and 4 months of rotenone administration, while 5.5 months of pesticide administration induced motor impairment as shown by the significant (p<0.05) decrease in force applied (Fig. 1D). However, there was no difference in latency to fall in the Rotarod test between control and rotenone-treated animals at any time point (Fig. 1C).

This confirmed that intranasal rotenone administration induced a slowly progressive motor dysfunction in mice after 5.5 months.

### Rotenone exposure induced atrophy of OB and aSyn accumulation

Next, the size of OB was evaluated from isolated brain hemispheres of vehicle and rotenone-treated animals. When measured by Vernier callipers, we found significant atrophy in OB of mice treated with rotenone for 4 and 5.5 months compared to controls. Rotenone exposure for 3 months did not induce any atrophy of OB (Fig. 2A).

**Figure 2.**
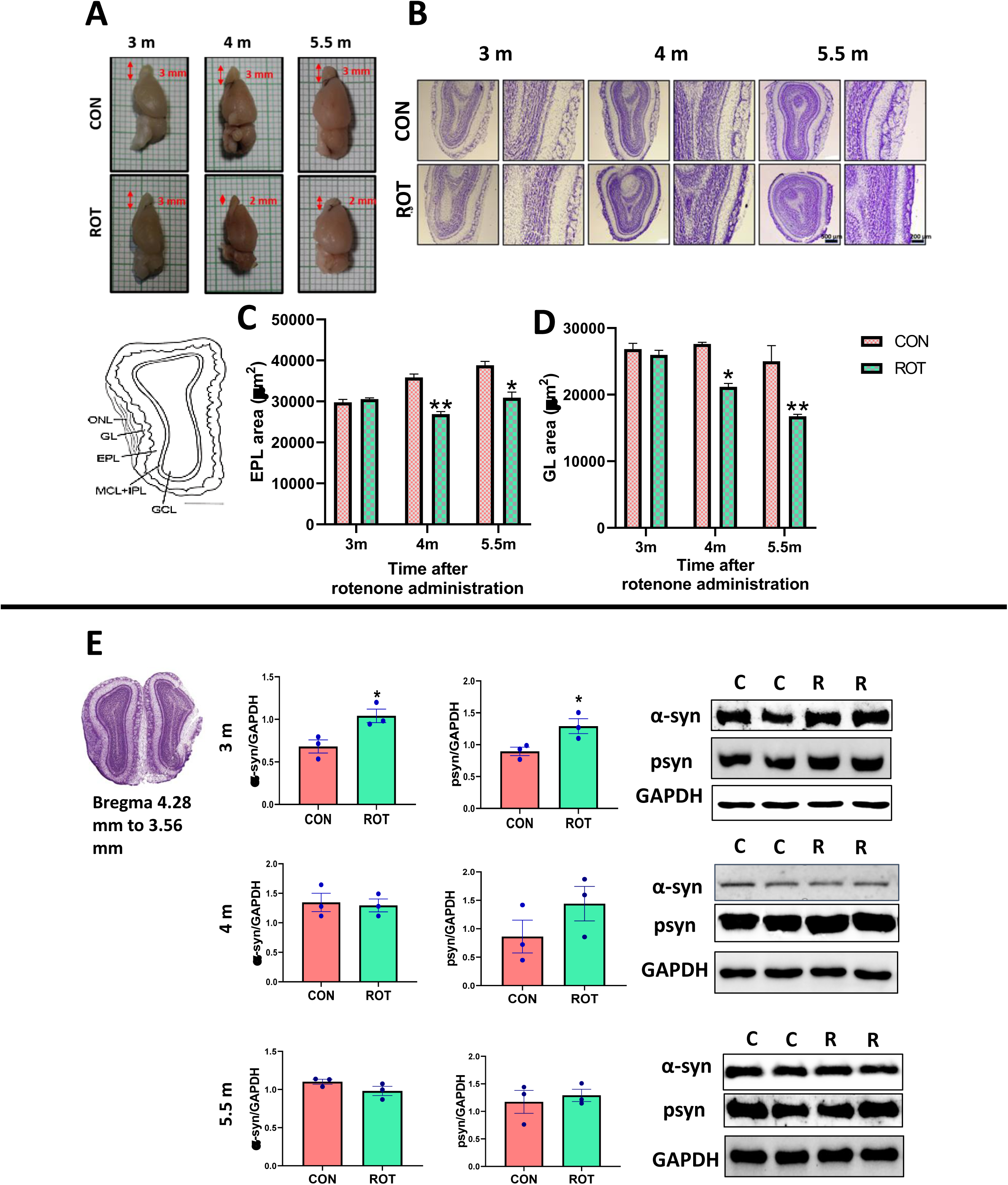
Intranasal rotenone induced atrophy of olfactory bulb (OB). A. Figure showing reduction in OB size after 4 and 5.5 months of rotenone administration. B. Nissl-stained coronal sections of OB at 3, 4 and 5.5 months after rotenone administration. C and D. Graphs represent the external plexiform layer (EPL) and glomerular layer (GL) areas at 3, 4 and 5.5 months after rotenone administration. Data was analysed by unpaired Student t-test and expressed as mean ± SEM (n=3), *p˂0.05, **p˂0.01 and ***p<0.001 vs control as observed after Nissl staining. E. Representative blots and quantification of α-syn and phosphorylated α-syn (psyn) in OB of control and rotenone animals after 3, 4 and 5.5 months of i.n. rotenone administration. Data was analyzed by unpaired t-test and expressed as mean ± SEM (n=3 in control group and n=3 in rotenone group), *p˂0.05 vs control

To further confirm the OB atrophy, we performed Nissl staining. We did not observe any atrophy in the coronal sections of OB after 3 months (Fig. 2B, 2C, 2D), while the area of the external plexiform layer was found to be significantly decreased in the rotenone group at 4 months (p<0.01) and 5.5 months’ (p<0.01) time points, along with the area of the glomerular layer, which was also significantly decreased after 4 (p<0.001) and 5.5 months (p<0.05) of rotenone exposure (Fig. 2C, 2D).

Western blot analysis showed a significant (p<0.05) increase in aSyn and pSyn expression in OB of rotenone-treated mice after 3 months of administration (p<0.05). There was no significant difference in aSyn expression in OB between control and rotenone-treated animals at 4 months and 5.5 months (Fig. 2E).

### Rotenone induced alpha synuclein accumulation in anatomically connected regions of the olfactory bulb

#### Anterior Olfactory Nucleus (AON)

IHC studies showed a significant increase in aSyn expression in AON after 3 months (p<0.05), 4 months (p<0.001) and 5.5 months (p<0.05) of rotenone administration (Fig. 3A) as compared to control. Similarly, immunofluorescence studies showed significantly increased pSyn expression in AON after 3 months (p<0.05), 4 months (p<0.01) and 5.5 months (p<0.05) of rotenone administration compared to control.(Fig. 4A).

**Figure 3.**
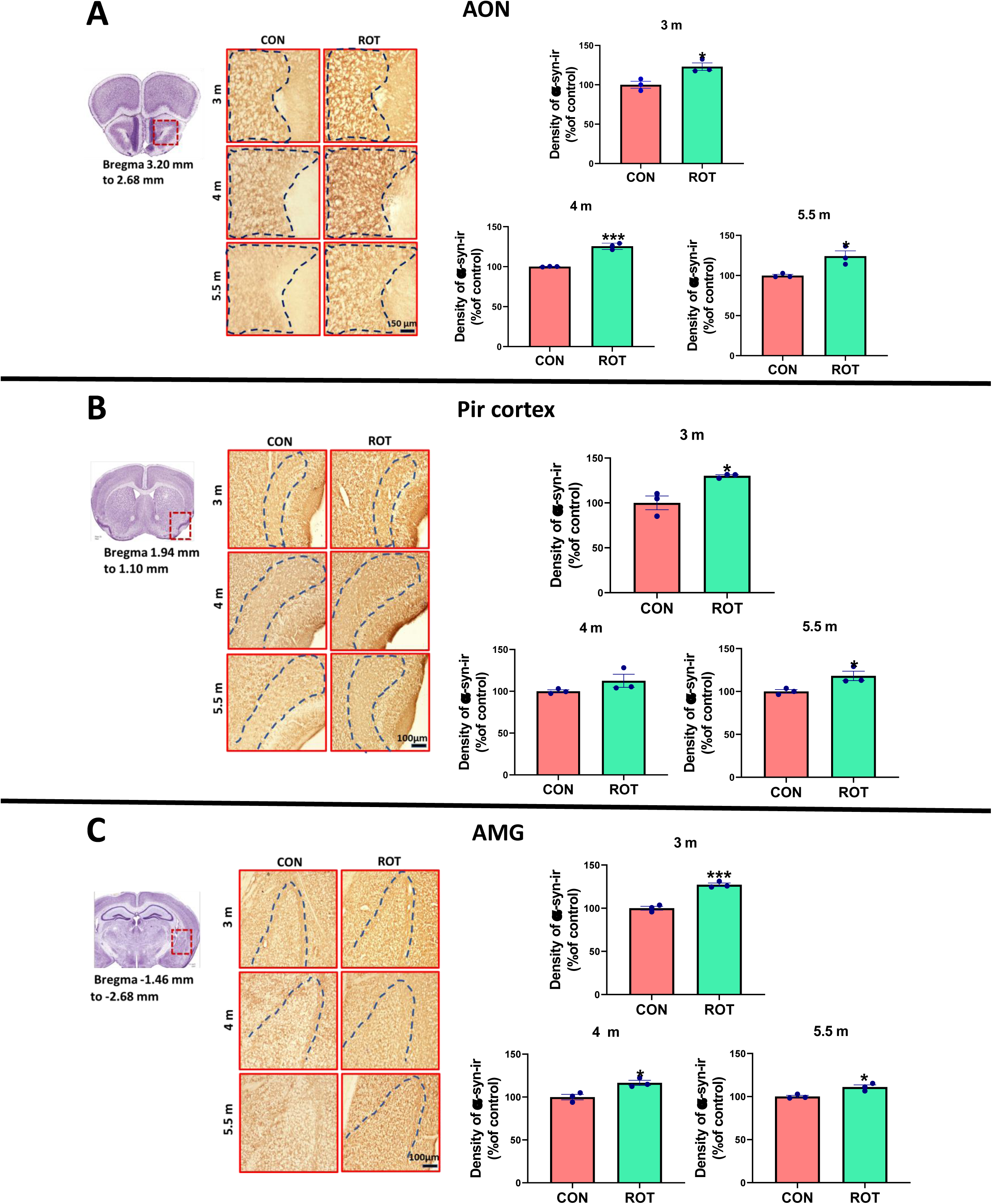
Rotenone induced aSyn accumulation in the anterior olfactory nucleus (AON), piriform cortex (Pir cortex) and amygdala (AMG). Figure represents the images of DAB-stained sections and graphs representing the α-syn immunoreactive density in (A) AON, (B) Pir cortex and (C) AMG of control and rotenone treated mice after 3,4 and 5.5 months of rotenone administration. Data was analyzed by unpaired t-test and expressed as mean ± SEM (n=3 in control group and n=3 in rotenone group), *p˂0.05 and ***p<0.001 vs control

**Fig 4.**
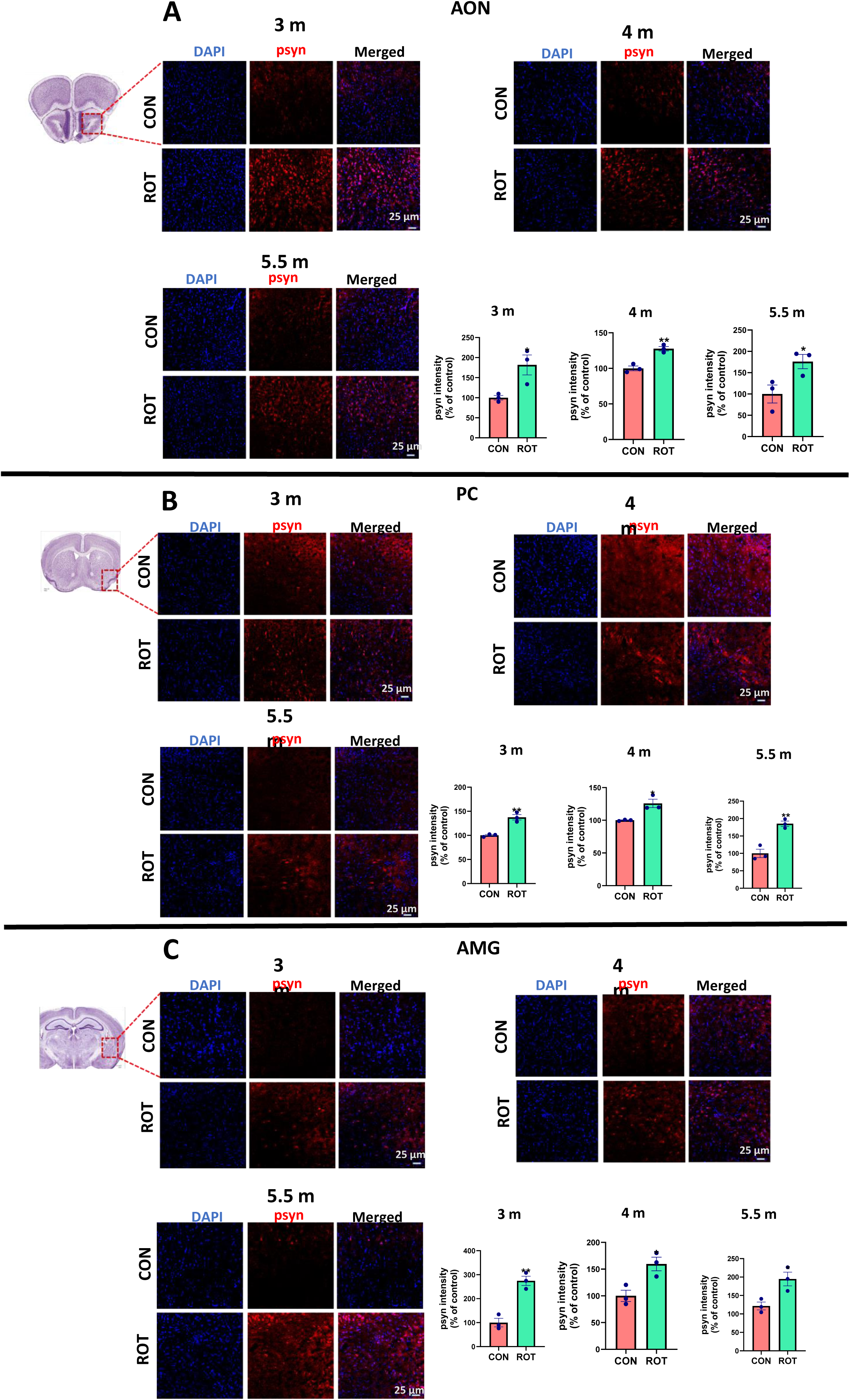
Rotenone enhanced pSyn expression in the anterior olfactory nucleus (AON), piriform cortex (Pir cortex) and amygdala (AMG). Representative images of immunofluorescence (IF) staining for co-localization of pSyn and DAPI in (A) AON, (B) Pir cortex and (C) AMG of mice at 40X magnification. Graphs represent the psyn intensity (% of control) in (A) AON, (B) Pir cortex and (C) AMG in both groups at 3, 4 and 5.5 months after rotenone administration. Data was analyzed by unpaired t-test and expressed as mean ± SEM (n=3 in control group and n=3 in rotenone group), *p˂0.05 and **p<0.01 vs control

#### Piriform cortex

IHC studies showed a significant increase in aSyn expression in piriform cortex after 3 months (p<0.05) and 5.5 months (p<0.05) of rotenone administration compared to control (Fig. 3B). psyn expression was also found to be significantly increased in piriform cortex after 3 months (p<0.01), 4 months (p<0.05) and 5.5 months (p<0.01) of rotenone administration (Fig. 4B).

#### Amygdala

IHC studies showed, significant increase in aSyn expression in amygdala after 3 months (p<0.001), 4 months (p<0.05) and 5.5 (p<0.05) months of rotenone administration as compared to control (Fig. 3C). With the help of immunofluorescence, psyn expression was found to be significantly increased in amygdala after 3 months (p<0.01), 4 months (p<0.05) and 5.5 (p<0.05) months of rotenone administration as compared to control (Fig. 4C).

### Rotenone induced alpha-synuclein accumulation in the nigrostriatal and cortical regions

#### Striatum and cortex

Western blot analysis showed aSyn pathology progression to striatum (p<0.05) (Fig. 5a) and cortex (p<0.05) (Fig. 5 B) only after 5.5 months of rotenone exposure. ***SN*:** With the help of immunofluorescence, we observed no difference in aSyn expression in SN of control and rotenone group animals after 3 and 4 months of administration, while aSyn intensity was found to be significantly (p<0.05) increased in SN of rotenone treated mice only after 5.5 months of administration (Fig. 6A). Analogous trend was observed in pSyn expression. Rotenone administration for 5.5 months caused a significant (p<0.05) increase in pSyn intensity as compared to control (Fig. 6B).

**Fig 5.**
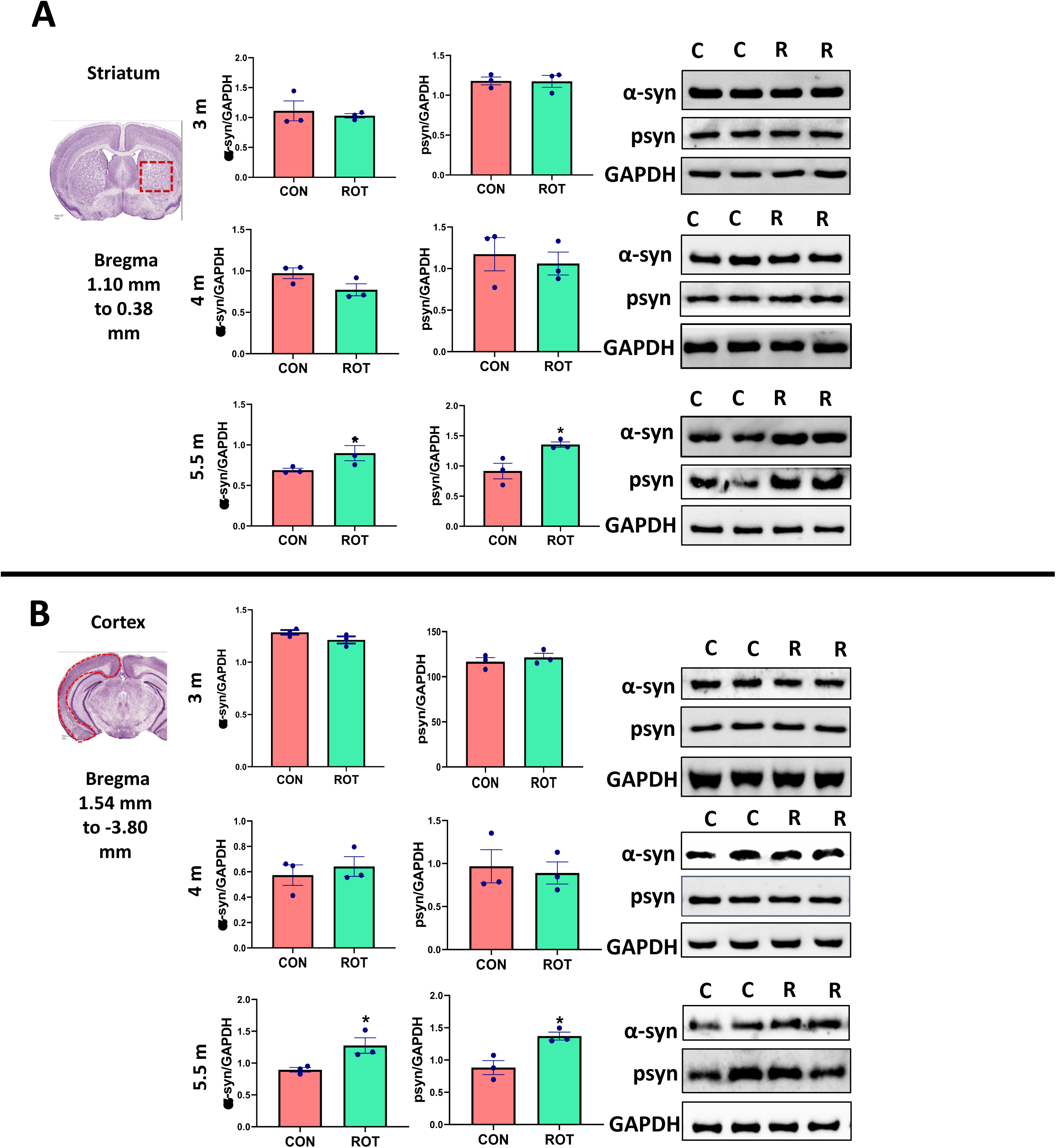
Rotenone induced aSyn accumulation in the straitum and cortex. Representative blots and quantification of α-syn and phosphorylated α-syn (psyn) in (A) striatum and (B) cortex of control and rotenone animals after 3, 4 and 5.5 months of intranasal rotenone ME administration: Data was analyzed by unpaired t-test and expressed as mean ± SEM (n=3 in control group and n=3 in rotenone group), *p˂0.05 vs control

**Fig 6.**
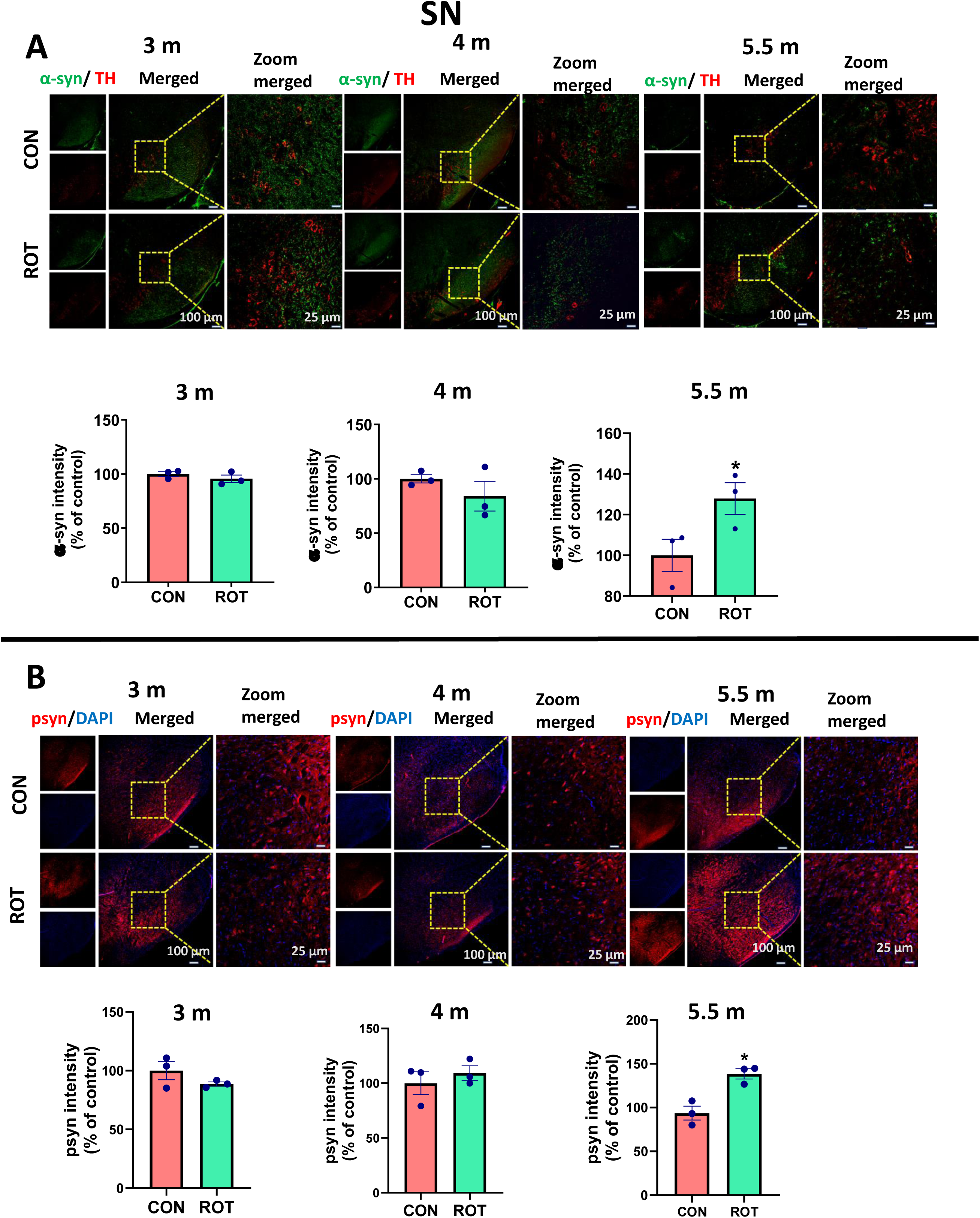
Rotenone induced aSyn aggregation in substantia nigra (SN). Representative images of immunofluorescence (IF) staining for co-localization of α-syn and TH expression in SN of mice at 10X and 40X magnification (A). Graphs representing the α-syn intensity (% of control) in TH+ neurons in SN in both groups at 3, 4 and 5.5 months after rotenone administration. Data was analyzed by unpaired t-test and expressed as mean ± SEM (n=3 in control group and n=3 in rotenone group), *p˂0.05 vs control (B) Rotenone administration for 3 months and 4 months didn’t change psyn expression, but for 5.5 months increased psyn intensity in substantia nigra: Representative images of immunofluorescence (IF) staining for co-localization of psyn and DAPI in SN of mice at 10X and 40X magnification. Graphs represent the psyn intensity (% of control) in SN in both groups at 3, 4 and 5.5 months after rotenone administration. Data was analyzed by unpaired t-test and expressed as mean ± SEM (n=3 in control group and n=3 in rotenone group), *p˂0.05 vs control

### Intranasal rotenone administration induced astroglial activation in the olfactory bulb and nigrostriatal regions

Next, we assessed astroglial cell activation in the OB, Striatum and SN.

#### OB

We observed a significant increase (p<0.05) in the GFAP expression marker for astroglial cell activation in OB after 3 months of rotenone administration, which subsided after 4 and 5.5 months of administration.

#### Striatum

In the striatum of rotenone-administered mice, there was significant astroglial activation at all three time points, 3 months (p<0.01), 4 months (p<0.05) and 5.5 months (p<0.05) (Fig. 8A).

*SN:* We found significant astroglial activation in SN after 4 (p<0.05) and 5.5 months (p<0.05), but not after 3 months (Fig. 9A and supplementary Fig.3).

### Intranasal rotenone administration induced alterations in phosphorylated tyrosine hydroxylase (pTH) levels in the olfactory bulb and striatum

Phosphorylated tyrosine hydroxylase, especially at serine 40, is known to significantly enhance TH activity, and it can be an earlier marker of dopaminergic neuronal loss. Hence, we determined its levels in these regions by Western blot analysis (Dunkley et al., 2004)

### OB

We investigated the levels of pTH using western blot and found decreased (p<0.05) pTH levels at all time points (Fig. 7A).

**Fig 7.**
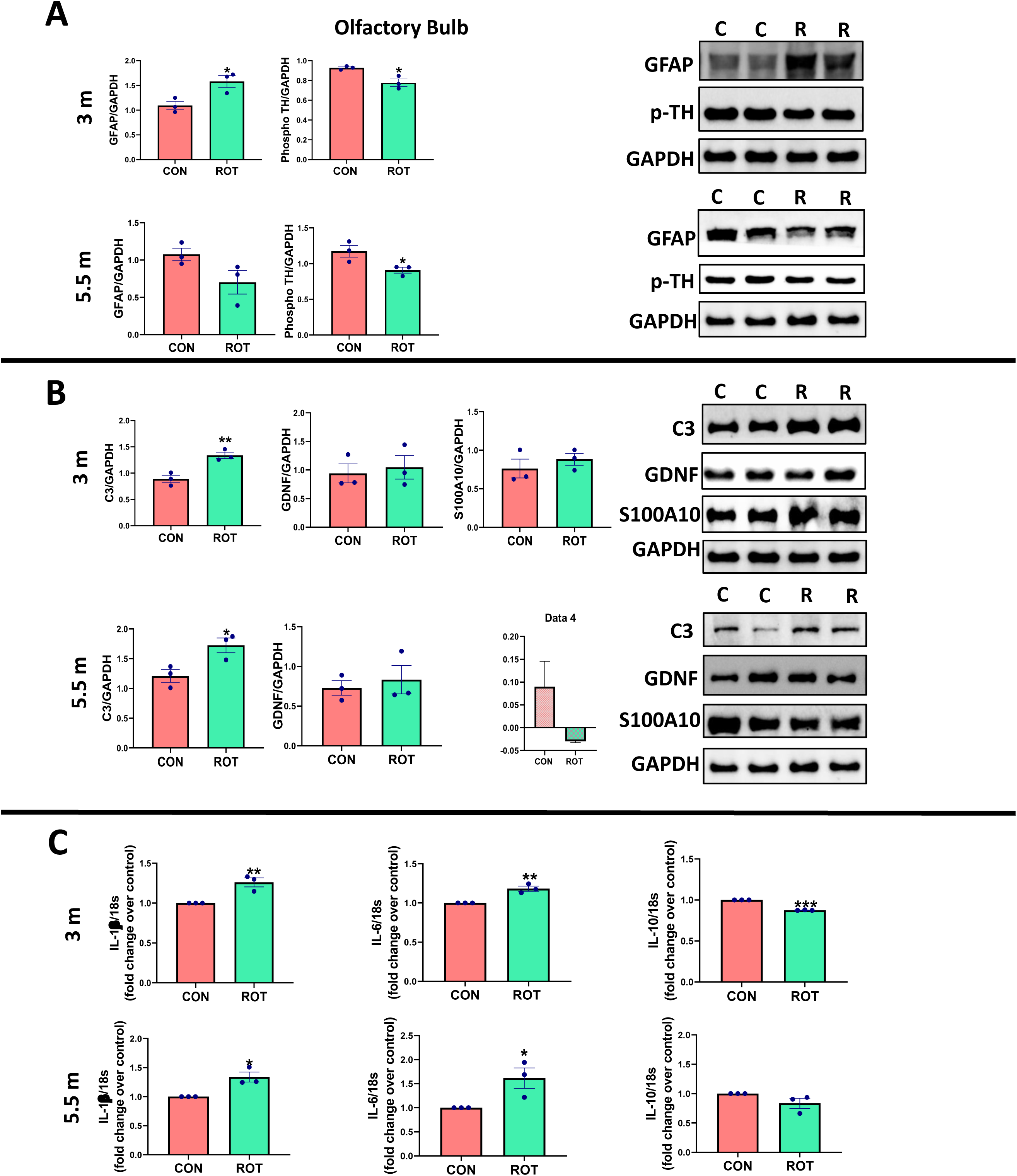
Rotenone induced differential reactive astrogliosis in the olfactory bulb (OB). (A) Representative blots and quantification of GFAP and pTH in the OB of control and rotenone animals after 3 and 5.5 months of intranasal rotenone ME administration. Data was analyzed by unpaired t-test and expressed as mean ± SEM (n=3 in control group and n=3 in rotenone group), *p˂0.05 vs control. (B) Representative blots and quantification of C3, GDNF and S100A10 in OB of control and rotenone animals after 3 and 5.5 months of intranasal rotenone ME administration. Data was analyzed by unpaired t-test and expressed as mean ± SEM (n=3 in control group and n=3 in rotenone group), *p˂0.05, **p˂0.01 vs control. (C) Rotenone enhanced the production of proinflammatory cytokines in OB after 3 as well as 5.5 months of administration. Rotenone decreased the anti-inflammatory cytokine production after 3 months, but induced no change in its expression after 5.5 months of administration. This figure represents the effect of rotenone on expression of IL-1β, IL-6 and IL-10 in OB after rotenone administration for 3 and 5.5 months. Data is expressed as mean±SEM (n=3). Data was analyzed by unpaired t-test and expressed as mean ± SEM (n=3 in control group and n=3 in rotenone group), *p<0.05, ***p<0.001 vs control

### Striatum

Importantly, we found an increase in pTH (p<0.05) levels at the 3-month time point using western blot. In contrast, it was found to be decreased (p<0.05) at the 5.5-month time point (Fig. 8A).

**Fig 8.**
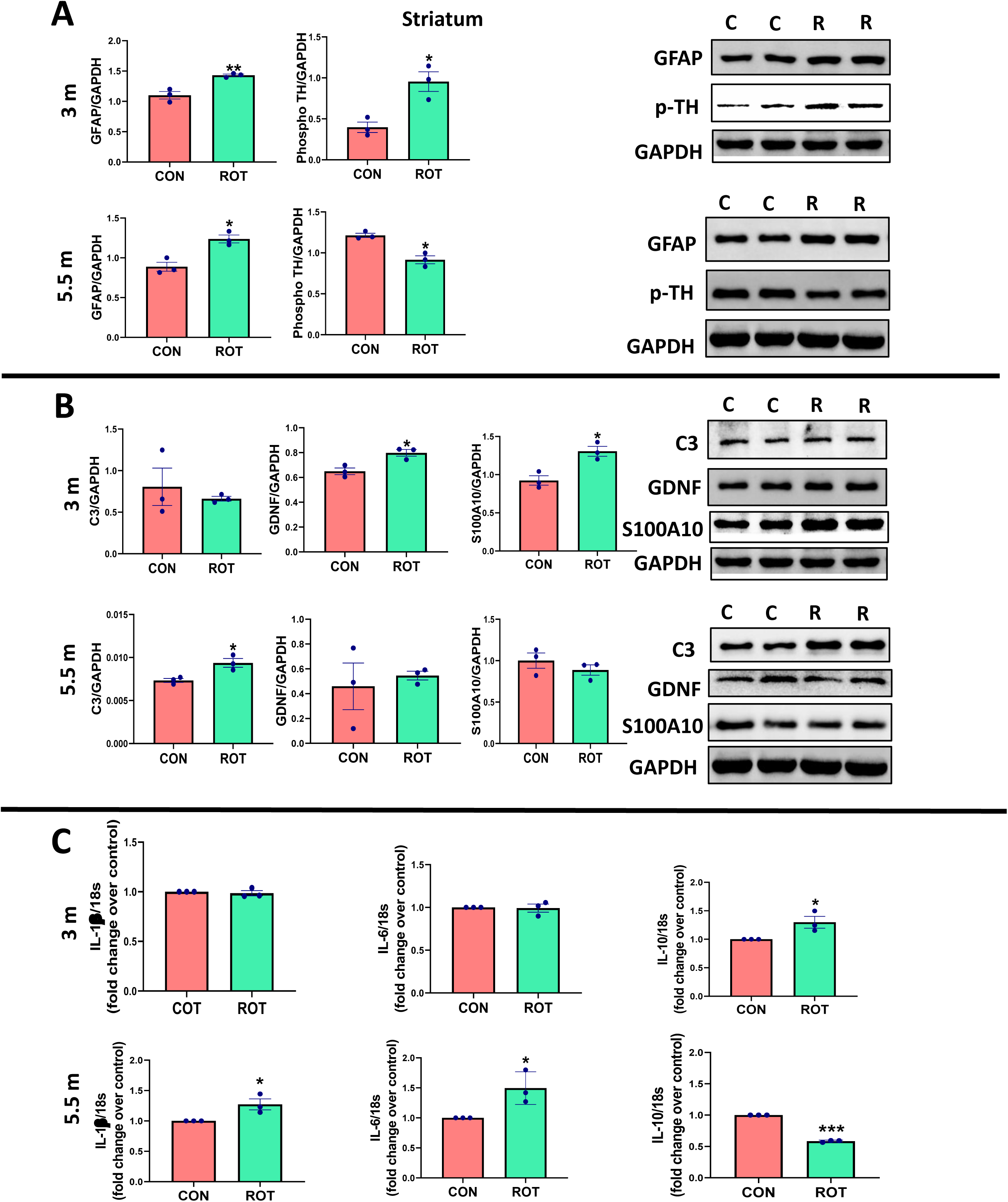
Rotenone induced differential reactive astrogliosis in the striatum. (A) Representative blots and quantification of GFAP and pTH in the striatum of control and rotenone animals after 3 and 5.5 months of intranasal rotenone ME administration. Data was analyzed by unpaired t-test and expressed as mean ± SEM (n=3 in control group and n=3 in rotenone group), *p˂0.05, **p˂0.01 vs control. (B) Representative blots and quantification of C3, GDNF and S100A10 in striatum of control and rotenone animals after 3 and 5.5 months of intranasal rotenone ME administration. Data was analyzed by unpaired t-test and expressed as mean ± SEM (n=3 in control group and n=3 in rotenone group), *p˂0.05 vs control. (C) Rotenone enhanced the production of proinflammatory cytokines in the striatum after 5.5 months of administration, but not after 3 months of administration. Rotenone decreased the anti-inflammatory cytokine production after 5.5 months, but increased its expression after 3 months of administration. This figure represents the effect of rotenone on the expression of IL-1β, IL-6 and IL-10 in the striatum after rotenone administration for 3 and 5.5 months. Data is expressed as mean±SEM (n=3). Data was analyzed by unpaired t-test and expressed as mean ± SEM (n=3 in control group and n=3 in rotenone group), *p<0.05, ***p<0.001 vs control

As we found a significant increase in GFAP expression in the striatum of rotenone-treated mice as compared to control mice after 3 months and 5.5 months, while we observed the opposite trend in expression of pTH (Fig. 8A) at these time points, we suspected the presence of different phenotypes of astroglia. Astrocytes may show different phenotypes based on the surrounding environment, A1 and A2 astrocytes. A1 astrocytes are supposed to be less supportive to neurons, whereas A2 astrocytes are protective or more supportive in nature (Ding *et al*. 2021a). The activation of A2 astrocytes is associated with an increase in GDNF expression (Li *et al*. 2019), further causing the increase in pTH levels. As the conversion of A2 astrocytes into A1 type occurs, it disrupts the normal function of astrocytes to release GDNF. We checked the expression of C3a (marker of A1 astrocytes) and S100A10 (marker of A2 astrocytes) in OB and striatum and the levels of pro-inflammatory and anti-inflammatory cytokines in the similar regions.

### Intranasal rotenone administration induced a time-dependent, distinct reactive phenotype of astroglia in OB, striatum and substantia nigra

#### OB

In OB, at 3 months, there was a significant (p<0.01) increase in expression of C3, while no change was observed in GDNF and S100A10 expression. At 5.5 months, expression of C3 was significantly (p<0.05) increased, with no change in GDNF expression and a significant (p<0.05) decrease in S100A10 expression (Fig. 7B).

There was a significant increase in expression of IL-1β and IL-6 in OB of rotenone-administered animals at 3 (p<0.01) as well as 5.5 months (p<0.05) time points, as compared to control. While expression of anti-inflammatory cytokine IL-10 was found to be significantly decreased (p<0.001) in OB of rotenone-administered animals at the 3-month time point (Fig. 7C), there was no change observed in expression of IL-10 after 5.5 months of rotenone administration as compared to control (Fig. 7C).

#### Striatum

We found a significant (p<0.05) increase in S100A10 expression marker for A1 astrocytes in rotenone-exposed mice, consistent with a significant (p<0.05) increase in GDNF expression as compared to control at the 3-month’ time point. (Fig. 8B).

There was a significant increase in expression of IL-1β in the striatum of rotenone-administered animals as compared to control at 5.5 months, while no change was observed at 3 months (Fig. 8C).

While expression of anti-inflammatory cytokine IL-10 was found to be significantly increased (p<0.001) in the striatum of rotenone-administered animals at the 3-month time point, there was a significant decrease (p<0.05) in expression of IL-10 after 5.5 months of rotenone administration as compared to control (Fig. 8C).

#### SN

Expression of C3 marker for A1 astrocytes was significantly (p<0.01) decreased in rotenone treated mice after 4 months of administration, while it was found to be increased (p<0.01) after 5.5 months as compared to control (Fig. 9C). Whereas expression of S100A10 marker for A2 astrocytes was significantly (p<0.05) increased (Fig. 9D) which was correlative with increase (p<0.05) in GDNF expression in rotenone group mice after 4 months (Fig. 9B), while there was no change observed after 5.5 months of rotenone exposure in S100A10 (Fig. 9D) with a significant (p<0.05) decrease in GDNF as compared to control (Fig. 9B).

**Fig 9.**
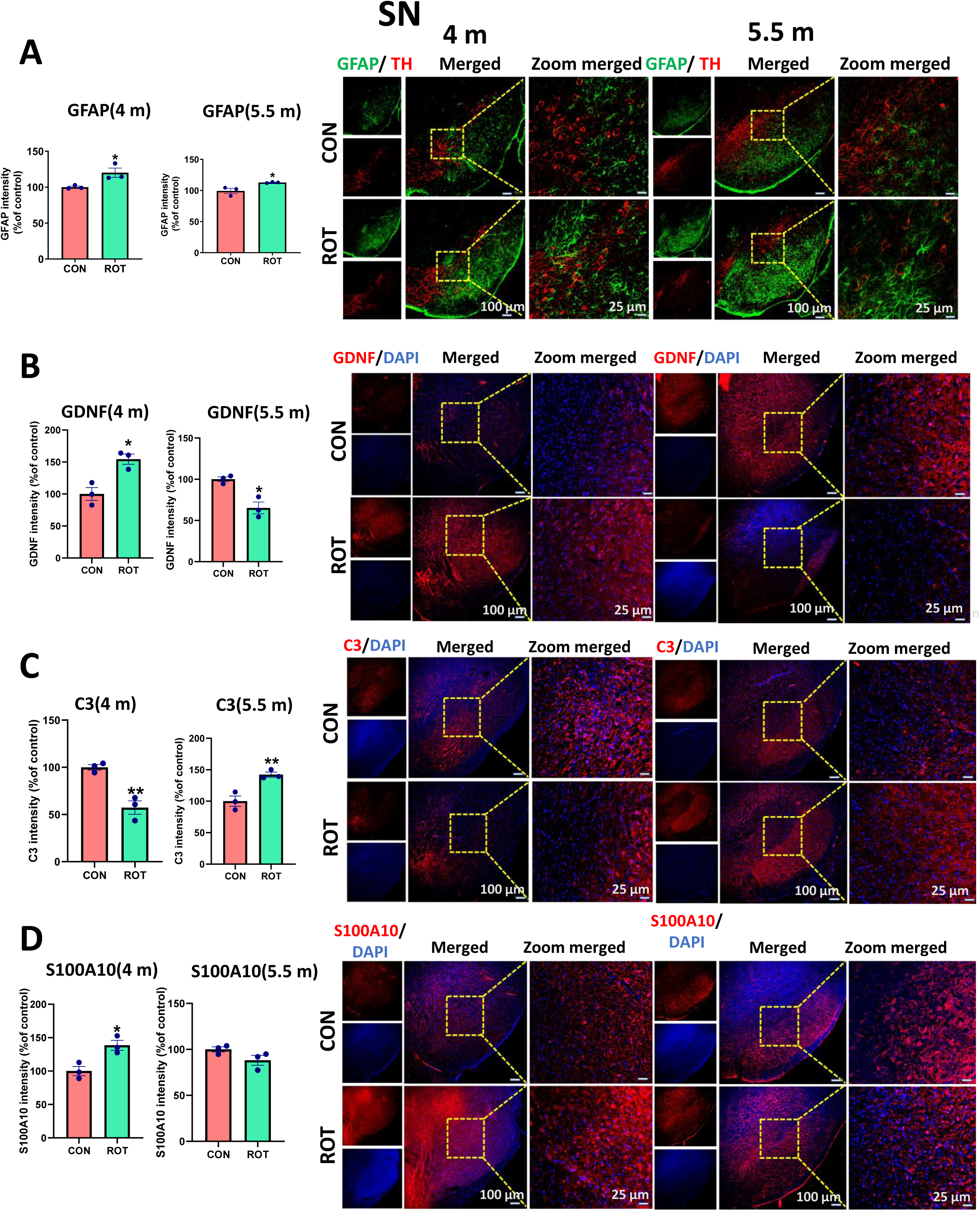
Rotenone induced differential reactive astrogliosis in the substantia nigra. (A) Rotenone increased GFAP expression in substantia nigra (SN) of mice after 4 and 5.5 months of intranasal administration, but didn’t change GFAP intensity after 3 months: Representative images of IF staining for co-localization of GFAP and TH expression in SN of mice at 10X and 40X magnification. Graph representing the GFAP intensity (% of control) in TH+ neurons in SN in both groups at 4 and 5.5 months after rotenone administration. Data was analyzed by unpaired t-test and expressed as mean ± SEM (n=3 in control group and n=3 in rotenone group), *p ˂0.05 vs control. (B) Rotenone increased C3 expression in SN of mice after 4 as well as 5.5 months of intranasal administration: Representative images of immunofluorescence (IF) staining for co-localization of C3 and TH expression in SN of mice at 10X and 40X magnification. Graph representing the C3 intensity (% of control) in TH+ neurons in SN in both groups at 4 and 5.5 months after rotenone administration. Data was analyzed by unpaired t-test and expressed as mean ± SEM (n=3 in control group and n=3 in rotenone group), **p ˂0.01 vs control. (C) Rotenone increased S100A10 expression in SN of mice after 4 months of intranasal administration, but didn’t change S100A10 intensity after 5.5 months: Representative images of immunofluorescence (IF) staining for co-localization of S100A10 and TH expression in substantia nigra of mice at 10X and 40X magnification. Graph representing the S100A10 intensity (% of control) in TH+ neurons in SN in both groups at 4 and 5.5 months after rotenone administration. Data was analyzed by unpaired t-test and expressed as mean ± SEM (n=3 in control group and n=3 in rotenone group), *p ˂0.05 vs control. (D) Intranasal administration of rotenone for 4 months increased the GDNF expression, but for 5.5 months, decreased the GDNF expression in SN of mice Representative images of immunofluorescence (IF) staining for co-localization of GDNF and TH expression in substantia nigra of mice at 10X and 40X magnification. Graph representing the GDNF intensity (% of control) in TH+ neurons in SN in both groups at 4 and 5.5 months after rotenone administration. Data was analyzed by unpaired t-test and expressed as mean ± SEM (n=3 in control group and n=3 in rotenone group), *p ˂0.05 vs control.

### Intranasal rotenone induced dopaminergic (DA) neurodegeneration in OB, striatum, cortex and SN

#### OB

Rotenone induced DAergic neurodegeneration in OB just after 3 months of administration, as indicated by significant decrease in TH expression in rotenone treated animals as compared to control, which was persistent even after 4 and 5.5 months (Fig. 10A). This is in continuation with the behavioural results, where we found olfactory dysfunction at all the three time points (Fig. 1B).

**Fig 10.**
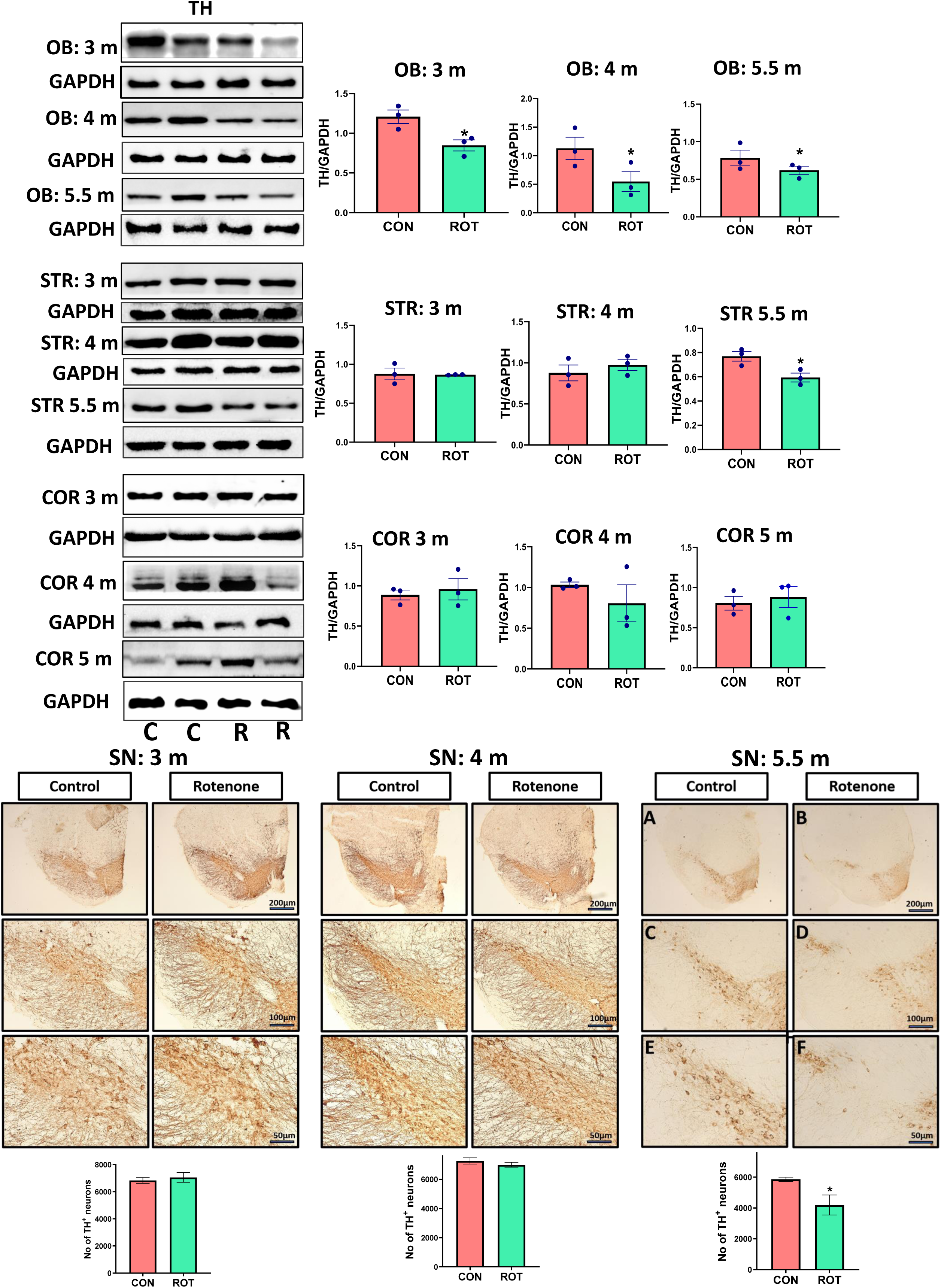
Rotenone induced dopaminergic neurodegeneration in olfactory bulb, striatum and substantia nigra. Representative blots and quantification of TH in OB, striatum and cortex of control and rotenone animals after 3, 4 and 5.5 months of intranasal rotenone ME administration (A, B and C). Data was analysed by unpaired t-test and expressed as mean ± SEM (n=3 in control group and n=3 in rotenone group), *p˂0.05 vs control. Figure represents the images of DAB-stained sections and graphs representing the number of dopaminergic neurons in the SN of control and rotenone-treated mice at 4x, 10x and 40x magnification after 3,4 and 5.5 months of rotenone administration (D). Data was analysed by unpaired t-test and expressed as mean ± SEM (n=3 in control group and n=3 in rotenone group), *p˂0.05 vs control

#### Striatum

We observed a significant decrease in TH expression in the striatum in rotenone-treated mice as compared to control after 5.5 months, which was not altered at 3- and 4-month (Fig. 10B) time points.

#### Cortex

We couldn’t see any changes in TH expression in cortex at any time point (Fig. 10C). ***SN***: There was no neurodegeneration observed in SN after 3 months and 4 months of rotenone administration, while 5.5 months of rotenone administration was able to induce significant (p ˂0.05) DAergic neurodegeneration in SN of rotenone-treated mice as compared to control (Fig. 10D).

## Discussion

The lack of an animal model of PD which can mimic the human PD condition hinders the development of drugs that can stop the progression of PD. It has been hypothesised that PD patients who show dysosmia or constipation a few decades before the development of motor symptoms may develop the aSyn pathology starting from the OB or gut, which later gets transferred to the midbrain. To mimic this phenomenon, an animal model that can simulate the disease progression from OB or gut to midbrain is required. Therefore, our study aimed to develop a model centred on the OB as the initial site of aSyn accumulation and investigate the evolution of pathology in other brain regions at different time points. As a result, we observed the time-dependent development of behavioural impairment and progression of aSyn pathology. In addition, we were able to analyse how neuroinflammation and neurodegeneration continue to undergo numerous changes over time and identify the most plausible relationship or mechanism underlying these alterations.

### Development and time-dependent progression of aSyn pathology from OB to other brain regions

Our findings from this rotenone model mimic the clinical situation. Olfactory impairment was observed just three months after rotenone administration, whereas motor dysfunction developed later. Due to the prodromal nature of this symptom, the olfactory test score can be a good diagnostic tool for assessing non-motor symptoms in PD, but it has limitations (Casjens *et al*. 2013). An olfactory deficit is common in PD, but it is not specific to the disease. The most commonly used screening tool for olfactory dysfunction is UPSIT, in which the patient is exposed to a specific fragrance, and it determines the individual’s smell function, which is mostly a qualitative response (Brumm M et. al., 2023). Moreover, it limits the ability to detect subtle differences in olfactory function (Vaswani *et al*. 2023). However, some researchers have observed a selective pattern of hyposmia in PD patients, suggesting the potential for developing better diagnostic tools (Double *et al*. 2003).

Braak in 2003 hypothesised the development of aSyn pathology from OB or enteric nervous system and its further progression to different brain regions via stereotypical progression based on post-mortem reports of PD patients (Braak *et al*. 2003). Rey et al. reported the progression from OB to 40 other brain regions after injecting different forms of aSyn in the olfactory bulb (Rey *et al*. 2013; Rey *et al*. 2016; Rey *et al*. 2018b). Similarly, we observed time-dependent progression of aSyn pathology from OB to the striatum, SN and piriform cortex via the olfactory pathway, though longitudinally. We confirmed the overexpression of aSyn in OB, AON, amygdala and piriform cortex of rotenone-treated mice after 3 months of rotenone administration, while this pathology reached the striatum, SN and cortex after 5.5 months. Neuronal projections (axons of mitral and tufted cells) emerging from the OB terminate in the olfactory cortex. The piriform cortex and anterior olfactory nucleus are also principal areas of the olfactory cortex (Haberly & Price 1978), along with the olfactory tubercle, anterior cortical amygdaloid nucleus, periamygdaloid cortex, and lateral entorhinal cortex (Ekanayake *et al*. 2023). That might be the reason for the simultaneous observation of aSyn pathology from OB to these areas. Rotenone causes *de novo* synthesis of aSyn (Sala *et al*. 2013). This aSyn accumulation in the initial stages might be the reason for the appearance of olfactory dysfunction observed after 3 months. Our results support the notion that aSyn can transfer along neural pathways and thereby contribute to the progression of the aSyn-related pathology.

### aSyn pathology subsides in atrophic OB after 4 and 5.5 months of rotenone administration

Surprisingly, there was no significant difference in aSyn expression in OB among control and rotenone animals at 4 and 5.5 months. One reason might be atrophy of OB; we found a significant decrease in the length and width of OBs in rotenone-treated mice. In rotenone-treated mice, distinct superficial layers of the OB, notably the olfactory nerve layer, glomerular layer, and external plexiform layer, shrank or shed considerably, resulting in atrophic OB. The mechanism behind this effect may be the loss of olfactory sensory neuron axons and injury to the olfactory epithelium, as shown before (Hasegawa-Ishii *et al*. 2019). However, this is beyond the scope of our research. Similarly, it has been stated that prolonged nasal irritation due to repeated exposure to lipopolysaccharide (LPS) is the factor responsible for atrophy (Hasegawa-Ishii et al. 2019). This is congruent with our results, in which repeated exposure of the nasal mucosa to rotenone led to persistent inflammation in OB, as evidenced by elevated GFAP levels in OB at all time points. In PD patients, clinical studies have also reported a decline in olfactory bulb volume, which is consistent with our findings (Li *et al*. 2016).

### Time-dependent differing role of astrocytes during the progression of aSyn pathology from OB to other brain regions

We found a variation in the expression of pTH in the striatum concerning GFAP expression at 3 and 5.5 months of rotenone administration. pTH levels were significantly higher at 3 months and significantly lower at 5.5 months in the striatum of the rotenone group, although GFAP expression was increased at both time points. An increase in GFAP expression represents reactive astrogliosis. Reactive astrogliosis can be a marker for the activation of A1 or A2 astrocytes (Liddelow & Barres 2017). A1 astrocytes generally get activated by neuroinflammation. They cause synaptic degeneration by activating genes in the complement cascade, which are known to damage synapses. Conversion of A2 to A1 leads to the release of neuroinflammatory factors and cytokines, including IL-1β, TNF-α, nitric oxide (NO), etc. Also loss of main functions of astrocytes, including synapse formation, phagocytosis of synapses, and support in neuronal survival and growth (Kim *et al*. 2020). A1 astrocytes release a toxin responsible for apoptosis of neurons and oligodendrocytes. A2 astrocytes generally get activated in ischemic conditions and are helpful as they upregulate many neurotrophic factors, that promote the survival and growth of neurons and thrombospondins, which aid in synapse repair (Liddelow & Barres 2017).

One of the growth factors released by A2 astrocytes is GDNF (glial-derived neurotrophic factor). GDNF acts as a potent survival factor for DAergic neurons and plays an important role in their phenotypic differentiation and maintenance. It prevents neuronal death caused by neurotoxin and axotomy and is also being seen as therapy for the treatment of PD (Patel & Gill 2007). Exposure to GDNF in neuroblastoma cells and primary neurons has been reported to increase TH phosphorylation at Ser-31 and Ser-40 positions, which enhances the activity of this enzyme and thus increases dopamine (DA) synthesis (Kobori *et al*. 2004). In the striatum, we observed activation of A2 astrocytes as revealed by increased levels of S100A10 (Li *et al*. 2020) at 3 months’ time point, which was not observed at 5.5 months. The activation of A2 astrocytes led to increase in GDNF expression, further causing the increase in pTH levels. We also found a decrease in proinflammatory cytokines, IL-6 and IL-1β, while anti-inflammatory cytokine (IL-10) levels were increased. Whereas at 5.5 months, A1 astrocytes were higher after rotenone exposure (as revealed by increase in expression of A1 marker complement C3 (Li *et al*. 2020)), leading to increased levels of IL-6 and IL-1β and decrease in levels of IL-10. There was no change in GDNF expression, consequently leading to a decrease in pTH levels in the rotenone group.

Existing literature suggests that activation of A1 astrocytes can contribute to neurodegeneration because of their abundance in neurodegenerative diseases (Li *et al*. 2019; Lawrence *et al*. 2023). This resembles our outcomes in SN, where we could observe loss of DAergic neurons with activation of A1 astrocytes at 5.5 months’ time point, which was not there after 4 months of administration. Transformation of astrocytes into A1 type is usually reported by release of IL-1α, tumor necrosis factor (TNF), and complement component 1q (C1q) by activated microglia (Liddelow *et al*. 2017), but in our study, we did not find any microglial activation, it might be possible that the microglial activation might happened prior to the 5.5 months time point or there might be other factors responsible in converting A2 astrocytes to A1 (Li *et al*. 2020). Similarly, A2 reactive astrocytes generally get induced by ischemia as reported in many studies (Gao *et al*. 2005; Hayakawa *et al*. 2014), and then they take part in CNS repair and recovery; however, studying the ischemic conditions in brain regions is beyond the scope of this publication. In our studies, the activation of A2 astrocytes might have improved neurogenesis at the 3-month’ time point.

### Time-dependent DAergic neurodegeneration from OB to other brain regions

We observed a decrease in TH expression in OB after 3 months of rotenone exposure, which was persistent even after 4 and 5.5 months. This aligns with our olfactory impairment results observed at all three time points. However, TH expression was found to be decreased in the striatum only after 5.5 months of administration. Similarly, we observed DAergic neuronal loss in SN after 5.5 months. This neuronal loss in SN might be due to aSyn accumulation or neuroinflammation and resulted in motor impairment observed in the grip strength test at this time point.

## Conclusion

Overall, we can conclude that we have developed a brain first chronic progressive mouse model of PD, which may mimic the development of pathology similar to PD patients and thus is suitable for testing any potential therapeutics for PD treatment. Most importantly, we were able to observe the protective role of astrocytes in the striatum and SN during the prodromal stage of the disease and it should be extensively studied in other animal models of PD. This opens the new avenue of future treatment with PD, as endogenous activation of astrocytes to A2 protective phenotype may protect the death of dopaminergic neurons. Future studies are warranted to study the autophagy, oxidative stress, neuroinflammation and other probable mechanisms involved in PD pathology progression. We suggest that similar studies can be repeated with older mice.

## Acknowledgements

The authors acknowledge National Institute of Pharmaceutical Education and Research (NIPER) Ahmedabad administration for providing the facility and support for conducting this study.

## Author contributions

All authors made fundamental contributions to the manuscript. AMK designed the study. MS, JS and MU conducted the experiments. MS analysed the data. MS, JS, JRK and AMK prepared the figures. AMK, JRK, JS, MU, MS, NS, TFO and IR participated in the interpretation and writing of the manuscript. All authors read and approved the final manuscript.

## Ethics approval

No clinical study was conducted. Experimental design for pre-clinical study was approved by IAEC Committee of NIPER Ahmedabad (IAEC approval number: NIPERA/IAEC/2021/010).

## References

Attems, J., Walker, L. and Jellinger, K. A. (2014) Olfactory bulb involvement in neurodegenerative diseases. Acta neuropathologica 127, 459–475.

Bandookwala, M., Sahu, A. K., Thakkar, D., Sharma, M., Khairnar, A., & Sengupta, P. (2019). Edaravone-caffeine combination for the effective management of rotenone induced Parkinson’s disease in rats: An evidence based affirmative from a comparative analysis of behavior and biomarker expression. Neuroscience Letters, 711, 134438. 10.1016/j.neulet.2019.134438

Berg, D., Postuma, R. B., Adler, C. H., Bloem, B. R., Chan, P., Dubois, B., … & Deuschl, G. (2015). MDS research criteria for prodromal Parkinson’s disease. Movement Disorders, 30(12), 1600–1611.

Bernis, M. E., Babila, J. T., Breid, S., Wüsten, K. A., Wüllner, U. and Tamgüney, G. (2015) Prion-like propagation of human brain-derived alpha-synuclein in transgenic mice expressing human wild-type alpha-synuclein. Acta neuropathologica communications 3, 1–18.

Borghammer, P., Horsager, J., Andersen, K., Van Den Berge, N., Raunio, A., Murayama, S., Parkkinen, L., & Myllykangas, L. (2021). Neuropathological evidence of body-first vs. brain-first Lewy body disease. Neurobiology of Disease, 161(November), 105557. 10.1016/j.nbd.2021.105557

Braak, H., Del Tredici, K., Rüb, U., De Vos, R. A., Steur, E. N. J. and Braak, E. (2003) Staging of brain pathology related to sporadic Parkinson’s disease. Neurobiology of aging 24, 197–211.

Casjens, S., Eckert, A., Woitalla, D. et al. (2013) Diagnostic value of the impairment of olfaction in Parkinson’s disease. PLoS One 8, e64735.

Ding, Z.-B., Song, L.-J., Wang, Q., Kumar, G., Yan, Y.-Q. and Ma, C.-G. (2021a) Astrocytes: a double-edged sword in neurodegenerative diseases. Neural Regeneration Research 16, 1702–1710.

Ding, Z.-B., Song, L.-J., Wang, Q., Kumar, G., Yan, Y.-Q. and Ma, C.-G. (2021b) Astrocytes: a double-edged sword in neurodegenerative diseases. Neural Regeneration Research 16, 1702.

Dunkley, P. R., Bobrovskaya, L., Graham, M. E., Von Nagy-Felsobuki, E. I., & Dickson, P. W. (2004). Tyrosine hydroxylase phosphorylation: regulation and consequences. Journal of Neurochemistry, 91(5), 1025–1043. 10.1111/j.1471-4159.2004.02797.x

Double, K. L., Rowe, D. B., Hayes, M. et al. (2003) Identifying the pattern of olfactory deficits in Parkinson disease using the brief smell identification test. Archives of neurology 60, 545–549.

Drolet, R. E., Cannon, J. R., Montero, L. and Greenamyre, J. T. (2009) Chronic rotenone exposure reproduces Parkinson’s disease gastrointestinal neuropathology. Neurobiology of disease 36, 96–102.

Ekanayake, A., Yang, Q., Kanekar, S., Ahmed, B., McCaslin, S., Kalra, D., Eslinger, P. and Karunanayaka, P. (2023) Monorhinal and Birhinal Odor Processing in Humans: an fMRI investigation. bioRxiv.

Farrand, A. Q., Verner, R. S., McGuire, R. M., Helke, K. L., Hinson, V. K. and Boger, H. A. (2020) Differential effects of vagus nerve stimulation paradigms guide clinical development for Parkinson’s disease. Brain Stimulation 13, 1323–1332.

Gao, Q., Li, Y. and Chopp, M. (2005) Bone marrow stromal cells increase astrocyte survival via upregulation of phosphoinositide 3-kinase/threonine protein kinase and mitogen-activated protein kinase kinase/extracellular signal-regulated kinase pathways and stimulate astrocyte trophic factor gene expression after anaerobic insult. Neuroscience 136, 123–134.

Haberly, L. B. and Price, J. L. (1978) Association and commissural fiber systems of the olfactory cortex of the rat. I. Systems originating in the piriform cortex and adjacent areas. Journal of Comparative Neurology 178, 711–740.

Hansen, C., Angot, E., Bergström, A.-L. et al. (2011) α-Synuclein propagates from mouse brain to grafted dopaminergic neurons and seeds aggregation in cultured human cells. The Journal of clinical investigation 121, 715–725.

Hasegawa-Ishii, S., Shimada, A. and Imamura, F. (2019) Neuroplastic changes in the olfactory bulb associated with nasal inflammation in mice. Journal of Allergy and Clinical Immunology 143, 978–989. e973.

Hayakawa, K., Pham, L.-D. D., Arai, K. and Lo, E. H. (2014) Reactive astrocytes promote adhesive interactions between brain endothelium and endothelial progenitor cells via HMGB1 and beta-2 integrin signaling. Stem cell research 12, 531–538.

Horsager, J., & Borghammer, P. (2024). Brain-first vs. body-first Parkinson’s disease: An update on recent evidence. Parkinsonism & Related Disorders, 122(January), 106101. 10.1016/j.parkreldis.2024.106101

Ip, C. W., Cheong, D. and Volkmann, J. (2017) Stereological estimation of dopaminergic neuron number in the mouse substantia nigra using the optical fractionator and standard microscopy equipment. *JoVE (Journal of Visualized Experiments)*, e56103.

Ishola, I., Awogbindin, I., Olubodun-Obadun, T., Olajiga, A. and Adeyemi, O. (2023) Vinpocetine prevents rotenone-induced Parkinson disease motor and non-motor symptoms through attenuation of oxidative stress, neuroinflammation and α-synuclein expressions in rats. Neurotoxicology 96, 37–52.

Jewett, M., Jimenez-Ferrer, I. and Swanberg, M. (2017) Astrocytic expression of GSTA4 is associated to dopaminergic neuroprotection in a rat 6-OHDA model of Parkinson’s disease. Brain sciences 7, 73.

Kawano, T. and Margolis, F. (1982) Transsynaptic regulation of olfactory bulb catecholamines in mice and rats. Journal of neurochemistry 39, 342–348.

Khairnar, A., Ruda-Kucerova, J., Drazanova, E., Szabó, N., Latta, P., Arab, A., Hutter-Paier, B., Havas, D., Windisch, M., Sulcova, A., Starcuk, Z., Király, A., & Rektorova, I. (2016). Late-stage α-synuclein accumulation in TNWT-61 mouse model of Parkinson’s disease detected by diffusion kurtosis imaging. Journal of Neurochemistry, 136(6), 1259–1269. 10.1111/jnc.13500

Khairnar, A., Plumitallo, A., Frau, L., Schintu, N. and Morelli, M. (2010) Caffeine enhances astroglia and microglia reactivity induced by 3, 4-methylenedioxymethamphetamine (‘ecstasy’) in mouse brain. Neurotoxicity research 17, 435–439.

Kim, E., Otgontenger, U., Jamsranjav, A. and Kim, S. S. (2020) Deleterious alteration of glia in the brain of alzheimer’s disease. International Journal of Molecular Sciences 21, 6676.

Kobori, N., Waymire, J. C., Haycock, J. W., Clifton, G. L. and Dash, P. K. (2004) Enhancement of tyrosine hydroxylase phosphorylation and activity by glial cell line-derived neurotrophic factor. Journal of Biological Chemistry 279, 2182–2191.

Kordower, Jeffrey H., Yaping Chu, Robert A. Hauser, Thomas B. Freeman, and C. Warren Olanow. “Lewy body–like pathology in long-term embryonic nigral transplants in Parkinson’s disease.” Nature medicine 14, no. 5 (2008): 504–506.

Lawrence, J. M., Schardien, K., Wigdahl, B. and Nonnemacher, M. R. (2023) Roles of neuropathology-associated reactive astrocytes: A systematic review. Acta Neuropathologica Communications 11, 1–28.

Li, Jia-Yi, Elisabet Englund, Janice L. Holton, Denis Soulet, Peter Hagell, Andrew J. Lees, Tammaryn Lashley et al. “Lewy bodies in grafted neurons in subjects with Parkinson’s disease suggest host-to-graft disease propagation.” Nature medicine 14, no. 5 (2008): 501–503.

Li, J., Gu, C.-z., Su, J.-b., Zhu, L.-h., Zhou, Y., Huang, H.-y. and Liu, C.-f. (2016) Changes in olfactory bulb volume in Parkinson’s disease: a systematic review and meta-analysis. PLoS One 11, e0149286.

Li, K., Li, J., Zheng, J. and Qin, S. (2019) Reactive astrocytes in neurodegenerative diseases. Aging and disease 10, 664.

Li, T., Liu, T., Chen, X., Li, L., Feng, M., Zhang, Y., Wan, L., Zhang, C. and Yao, W. (2020) Microglia induce the transformation of A1/A2 reactive astrocytes via the CXCR7/PI3K/Akt pathway in chronic post-surgical pain. Journal of Neuroinflammation 17, 1–15.

Liddelow, S. A. and Barres, B. A. (2017) Reactive astrocytes: production, function, and therapeutic potential. Immunity 46, 957–967.

Liddelow, S. A., Guttenplan, K. A., Clarke, L. E. et al. (2017) Neurotoxic reactive astrocytes are induced by activated microglia. Nature 541, 481–487.

Lin, L.-F. H., Doherty, D. H., Lile, J. D., Bektesh, S. and Collins, F. (1993) GDNF: a glial cell line-derived neurotrophic factor for midbrain dopaminergic neurons. Science 260, 1130–1132.

Luk, K. C., Song, C., O’Brien, P., Stieber, A., Branch, J. R., Brunden, K. R., Trojanowski, J. Q. and Lee, V. M.-Y. (2009) Exogenous α-synuclein fibrils seed the formation of Lewy body-like intracellular inclusions in cultured cells. Proceedings of the National Academy of Sciences 106, 20051–20056.

Miyazaki, I. and Asanuma, M. (2020) Neuron-astrocyte interactions in Parkinson’s disease. Cells 9, 2623.

Mougenot, A.-L., Nicot, S., Bencsik, A., Morignat, E., Verchère, J., Lakhdar, L., Legastelois, S. and Baron, T. (2012) Prion-like acceleration of a synucleinopathy in a transgenic mouse model. Neurobiology of aging 33, 2225–2228.

Parkhe, A., Parekh, P., Nalla, L. V., Sharma, N., Sharma, M., Gadepalli, A., Kate, A. and Khairnar, A. (2020) Protective effect of alpha mangostin on rotenone induced toxicity in rat model of Parkinson’s disease. Neuroscience Letters 716, 134652.

Patel, N. and Gill, S. S. (2007) GDNF delivery for Parkinson’s disease. Operative Neuromodulation, 135-154.

Prediger, R. D., Aguiar, A. S., Matheus, F. C., Walz, R., Antoury, L., Raisman-Vozari, R. and Doty, R. L. (2012) Intranasal administration of neurotoxicants in animals: support for the olfactory vector hypothesis of Parkinson’s disease. Neurotoxicity research 21, 90–116.

Rey, N. L., George, S., Steiner, J. A., Madaj, Z., Luk, K. C., Trojanowski, J. Q., Lee, V. M.-Y. and Brundin, P. (2018a) Spread of aggregates after olfactory bulb injection of α-synuclein fibrils is associated with early neuronal loss and is reduced long term. Acta neuropathologica 135, 65–83.

Rey, N. L., Petit, G. H., Bousset, L., Melki, R. and Brundin, P. (2013) Transfer of human α-synuclein from the olfactory bulb to interconnected brain regions in mice. Acta neuropathologica 126, 555–573.

Rey, N. L., Steiner, J. A., Maroof, N., Luk, K. C., Madaj, Z., Trojanowski, J. Q., Lee, V. M.-Y. and Brundin, P. (2016) Widespread transneuronal propagation of α-synucleinopathy triggered in olfactory bulb mimics prodromal Parkinson’s disease. Journal of Experimental Medicine 213, 1759–1778.

Rey, N. L., Wesson, D. W. and Brundin, P. (2018b) The olfactory bulb as the entry site for prion-like propagation in neurodegenerative diseases. Neurobiology of disease 109, 226–248.

Rojo, A. I., Cavada, C., de Sagarra, M. R. and Cuadrado, A. (2007) Chronic inhalation of rotenone or paraquat does not induce Parkinson’s disease symptoms in mice or rats. Experimental Neurology 208, 120–126.

Sala, G., Arosio, A., Stefanoni, G., Melchionda, L., Riva, C., Marinig, D., Brighina, L. and Ferrarese, C. (2013) Rotenone upregulates alpha-synuclein and myocyte enhancer factor 2D independently from lysosomal degradation inhibition. BioMed research international 2013.

Sasajima, H., Miyazono, S., Noguchi, T. and Kashiwayanagi, M. (2015) Intranasal administration of rotenone in mice attenuated olfactory functions through the lesion of dopaminergic neurons in the olfactory bulb. Neurotoxicology 51, 106–115.

Sasajima, H., Miyazono, S., Noguchi, T. and Kashiwayanagi, M. (2017) Intranasal administration of rotenone to mice induces dopaminergic neurite degeneration of dopaminergic neurons in the substantia nigra. Biological and Pharmaceutical Bulletin 40, 108–112.

Sharma, M., Malim, F. M., Goswami, A., Sharma, N., Juvvalapalli, S. S., Chatterjee, S., Kate, A. S., & Khairnar, A. (2023). Neuroprotective Effect of Swertiamarin in a Rotenone Model of Parkinson’s Disease: Role of Neuroinflammation and Alpha-Synuclein Accumulation. ACS Pharmacology & Translational Science, 6(1), 40–51. 10.1021/acsptsci.2c00120

Sharma, N., Soni, R., Sharma, M., Chatterjee, S., Parihar, N., Mukarram, M., Kale, R., Sayyed, A. A., Behera, S. K., & Khairnar, A. (2022). Chlorogenic Acid: a Polyphenol from Coffee Rendered Neuroprotection Against Rotenone-Induced Parkinson’s Disease by GLP-1 Secretion. Molecular Neurobiology, 59(11), 6834–6856. 10.1007/s12035-022-03005-z

Sharma, M., Kaur, J., Rakshe, S., Sharma, N., Khunt, D. and Khairnar, A. (2022) Intranasal Exposure to Low-Dose Rotenone Induced Alpha-Synuclein Accumulation and Parkinson’s Like Symptoms Without Loss of Dopaminergic Neurons. Neurotoxicity research 40, 215–229.

Sharma, M., Sharma, N. and Khairnar, A. (2023) Intranasal Rotenone Induces Alpha-Synuclein Accumulation, Neuroinflammation and Dopaminergic Neurodegeneration in Middle-Aged Mice. Neurochemical Research 48, 1543–1560.

Shiotsuki, H., Yoshimi, K., Shimo, Y., Funayama, M., Takamatsu, Y., Ikeda, K., Takahashi, R., Kitazawa, S. and Hattori, N. (2010) A rotarod test for evaluation of motor skill learning. Journal of neuroscience methods 189, 180–185.

Tepper, B., Koguc-Sobolewska, P., Jaslan, K., Turlejski, K., Bartkowska, K. and Djavadian, R. (2021) Impaired olfactory neurogenesis affects the performance of olfactory-guided behavior in aged female opossums. Scientific reports 11, 1–11.

Torres-Pasillas, G., Chi-Castañeda, D., Carrillo-Castilla, P., Marín, G., Hernández-Aguilar, M. E., Aranda-Abreu, G. E., Manzo, J. and García, L. I. (2023) Olfactory Dysfunction in Parkinson’s Disease, Its Functional and Neuroanatomical Correlates. NeuroSci 4, 134–151.

Vaswani, P. A., Morley, J. F., Jennings, D., Siderowf, A. and Marek, K. (2023) Predictive value of abbreviated olfactory tests in prodromal Parkinson disease. npj Parkinson’s Disease 9, 103.

Volpicelli-Daley, L. A., Luk, K. C., Patel, T. P., Tanik, S. A., Riddle, D. M., Stieber, A., Meaney, D. F., Trojanowski, J. Q. and Lee, V. M.-Y. (2011) Exogenous α-synuclein fibrils induce Lewy body pathology leading to synaptic dysfunction and neuron death. Neuron 72, 57–71.

Voronkov, D., Kutukova, K., Ivanov, M. and Khudoerkov, R. (2017) Immunomorphological changes in the olfactory bulbs of rats after intranasal administration of rotenone. Bulletin of experimental biology and medicine 164, 203–206.

Wang, S.-j., Wang, Q., Ma, J., Yu, P.-h., Wang, Z.-m. and Wang, B. (2018) Effect of moxibustion on mTOR-mediated autophagy in rotenone-induced Parkinson’s disease model rats. Neural regeneration research 13, 112.

Xavier, L. L., Viola, G. G., Ferraz, A. C., Da Cunha, C., Deonizio, J. M. D., Netto, C. A. and Achaval, M. (2005) A simple and fast densitometric method for the analysis of tyrosine hydroxylase immunoreactivity in the substantia nigra pars compacta and in the ventral tegmental area. Brain Research Protocols 16, 58–64.

