## Supplementary Data for "A ‘brain-first’ mouse model of progressive alpha-synuclein pathology via intranasal rotenone administration"

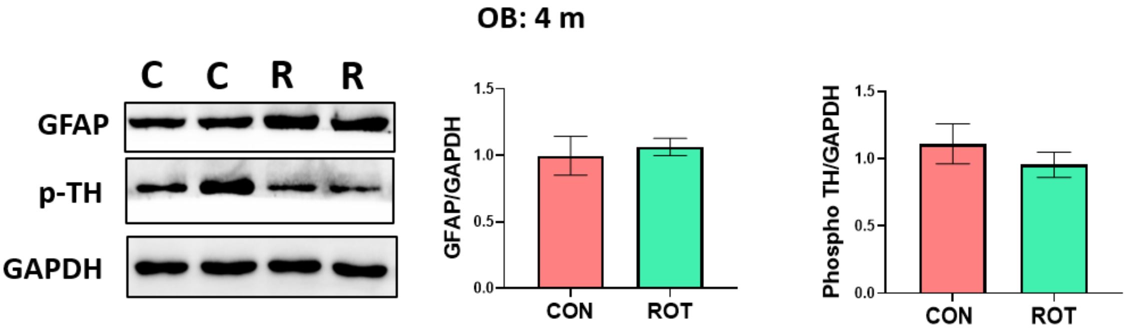


**Suppl. Fig. 1** Representative blots and quantification of GFAP and pTH in olfactory bulb (OB) of control and rotenone animals after 4 months of intranasal rotenone ME administration. Data was analyzed by unpaired t-test and expressed as mean ± SEM (n=3 in control group and n=3 in rotenone group).

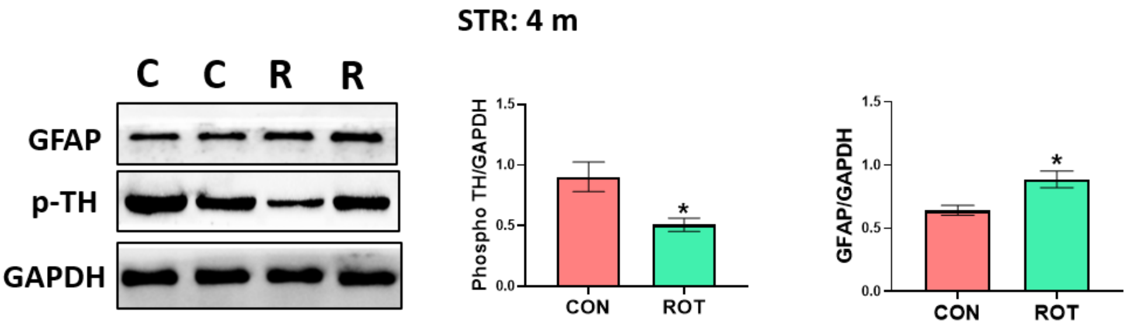


**Suppl. Fig. 2** Representative blots and quantification of GFAP and pTH in striatum of control and rotenone animals after 4 months of intranasal rotenone ME administration. Data was analyzed by unpaired t-test and expressed as mean ± SEM (n=3 in control group and n=3 in rotenone group), *p<0.05 vs control.

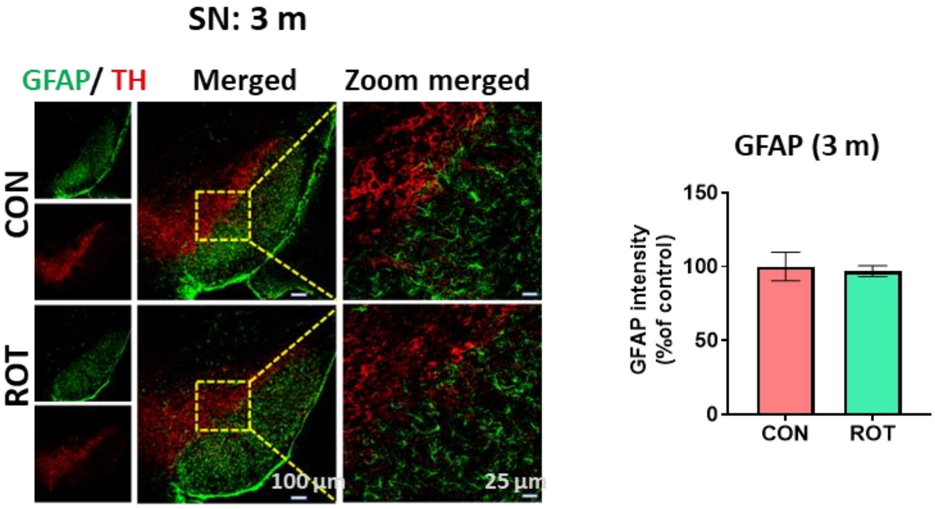


**Suppl. Fig. 3** Representative images of IF staining for co-localization of GFAP and TH expression in SN of mice at 10X and 40X magnification. Graph representing the GFAP intensity (% of control) in TH+

neurons in SN in both groups at 3 months after rotenone administration. Data was analyzed by unpaired t-test and expressed as mean ± SEM (n=3 in control group and n=3 in rotenone group)
